# Combining genome-wide studies of breast, prostate, ovarian and endometrial cancers maps cross-cancer susceptibility loci and identifies new genetic associations

**DOI:** 10.1101/2020.06.16.146803

**Authors:** Siddhartha P. Kar, Sara Lindström, Rayjean J. Hung, Kate Lawrenson, Marjanka K. Schmidt, Tracy A. O’Mara, Dylan M. Glubb, Jonathan P. Tyrer, Joellen M. Schildkraut, Jenny Chang-Claude, Ahmad G. M. Alsulimani, Fernando M. Antón, Alicia Beeghly-Fadiel, Line Bjørge, Clara Bodelon, Hiltrud Brauch, Stefanie Burghaus, Daniele Campa, Michael Carney, Chu Chen, Zhihua Chen, Mary B. Daly, Andreas du Bois, Arif B. Ekici, Ailith Ewing, Peter Fasching, James M. Flanagan, Jan Gawelko, Graham G. Giles, Robert J. Hamilton, Holly R. Harris, Florian Heitz, Michelle Hildebrandt, Peter Hillemanns, Ruea-Yea Huang, Liher Imaz, Arvids Irmejs, Anna Jakubowska, Allan Jensen, Esther M. John, Päivi Kannisto, Beth Y. Karlan, Elza Khusnutdinova, Lambertus A. Kiemeney, Susanne K. Kjaer, Rüdiger Klapdor, Petra Kleiblova, Martin Köbel, Bozena Konopka, Camilla Krakstad, Davor Lessel, Artitaya Lophatananon, Taymaa May, Agnieszka D. Mieszkowska, Alvaro N. Monteiro, Kirsten Moysich, Kenneth Muir, Sune F. Nielsen, Kunle Odunsi, Håkan Olsson, Tjoung-Won Park-Simon, Jennifer B. Permuth, Paolo Peterlongo, Agnieszka Podgorski, Ross Prentice, Paolo Radice, Harvey A. Risch, Ingo B. Runnebaum, Iwona K. Rzepecka, Rodney J. Scott, Veronica W. Setiawan, Nadeem Siddiqui, Weiva Sieh, Beata Śpiewankiewicz, Lukasz M. Szafron, Cheryl L. Thompson, Linda J. Titus, Clare Turnbull, Nawaid Usmani, Anne M. van Altena, Ana Vega-Gliemmo, Ignace Vergote, Robert A. Vierkant, Joseph Vijai, Stacey J. Winham, Robert Winqvist, Herbert Yu, the PRACTICAL consortium, CRUK, BPC3, CAPS, PEGASUS, Diether Lambrechts, Deborah J. Thompson, Ellen L. Goode, Wei Zheng, Ian P. M. Tomlinson, Andrew Berchuck, Susan J. Ramus, Stephen J. Chanock, Douglas F. Easton, Georgia Chenevix-Trench, Simon A. Gayther, Amanda B. Spurdle, Rosalind A. Eeles, Peter Kraft, Paul D. P. Pharoah

## Abstract

We report a meta-analysis of breast, prostate, ovarian, and endometrial cancer genome-wide association data (effective sample size: 237,483 cases/317,006 controls). This identified 465 independent lead variants (*P*<5×10^−8^) across 192 genomic regions. Four lead variants were >1Mb from previously identified risk loci for the four cancers and an additional 23 lead variant-cancer associations were novel for one of the cancers. Bayesian models supported pleiotropic effects involving at least two cancers at 222/465 lead variants in 118/192 regions. Gene-level association analysis identified 13 shared susceptibility genes (*P*<2.6×10^−6^) in 13 regions not previously implicated in any of the four cancers and not uncovered by our variant-level meta-analysis. Several lead variants had opposite effects across cancers, including a cluster of such variants in the TP53 pathway. Fifty-four lead variants were associated with blood cell traits and suggested genetic overlaps with clonal hematopoiesis. Our study highlights the remarkable pervasiveness of pleiotropy across hormone-related cancers, further illuminating their shared genetic and mechanistic origins at variant- and gene-level resolution.

## INTRODUCTION

Cancers of the breast, prostate, ovary and endometrium together accounted for nearly 22% of all new cancer cases diagnosed and approximately 1.2 million deaths worldwide in 2018^1^. These four cancers share common etiologies. Disease risks have been linked to variations in sex steroid hormones (e.g., hormone replacement therapy increases the risks of breast, ovarian, and endometrial cancers^2–4^) and shared genetic risk factors (pleiotropy). For example, rare highly penetrant germline mutations in the *BRCA1* and *BRCA2* genes elevate risks of breast, ovarian, and prostate cancers and have yielded fundamental cross-cancer mechanistic and therapeutic insights^5^. This provides the motivation for the identification of additional shared genetic risk factors, including pleiotropic common variants that may affect the risk of multiple cancers.

In 2016, we combined summary data from individual genome-wide association studies (GWAS) of breast, prostate, and ovarian cancer susceptibility that included a total of 112,349 cancer cases and 116,421 controls to identify 12 new cancer risk loci^6^. Since then, most of these loci have been replicated at genome-wide significance (*P* < 5 × 10^−8^) in larger, single-cancer GWAS of breast^7^, prostate^8^, or ovarian cancer^9^. We have now combined the results from these larger GWAS together with the results from an endometrial cancer GWAS^10^ in order to address four main aims. First, to combine single nucleotide polymorphism (SNP)-level associations across breast, prostate, ovarian, and endometrial cancers to identify susceptibility loci that have not been previously reported for any of the four cancers. Second, to use the combined data set to identify novel susceptibility loci for one of the four cancers in genomic regions that contain a known susceptibility locus for another of the four cancers. Third, to determine the combination of the four cancers that is most likely to underlie the association at the lead SNPs of all susceptibility loci identified in the combined data set in order to map shared or cross-cancer loci. Fourth, to carry out gene-level association analysis based on this powerful combined SNP-level data set to identify candidate cancer susceptibility genes in loci not previously identified by single- and multi-cancer SNP-level association analyses.

## RESULTS

### Cross-cancer genome-wide association meta-analysis

We used meta-analysis based on the Han and Eskin model^11,12^ to combine summary results for 9,530,997 variants with minor allele frequency > 1% from the largest genome-wide association data sets for susceptibility to breast^7^, prostate^8^, ovarian^9^, and endometrial cancers^10^ published as of August 2019 (**Methods**). All cases and controls included were of European ancestry (**Supplementary Table 1**). A recent study^13^ has shown that some breast cancer susceptibility SNPs have subtype-specific associations where the allele that confers risk of developing one subtype is protective for another subtype. Due to their opposite directions, such allelic associations may be masked in an analysis that does not take into account subtype-specific associations. Subtype-specific data were available for the current study by estrogen receptor (ER)-status for breast cancer and for high-grade serous ovarian cancer (HGSOC), which is the most common histological subtype of epithelial ovarian cancer and known to be similar at the molecular level with ER-negative breast cancer^14^. Therefore, we also conducted an additional meta-analysis combining the results for ER-positive breast cancer, ER-negative breast cancer, HGSOC, prostate cancer, and endometrial cancer. Hereafter, we refer to this additional analysis as the subtype-focused meta-analysis.

**Table 1.**
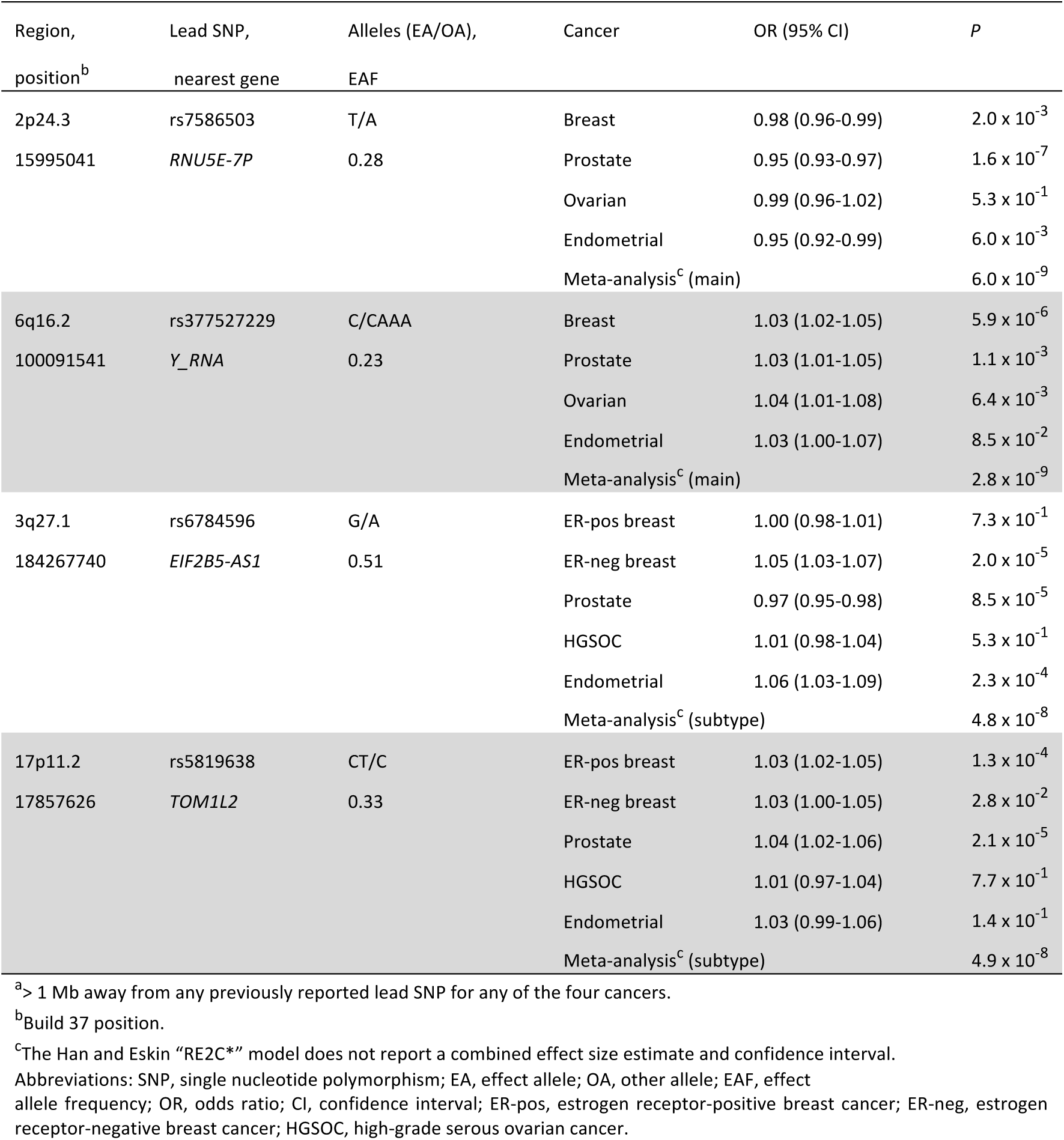
The cancer susceptibility loci not previously identified for any of the four cancers.^a^

| Region,<br>position<sup>b</sup> | Lead SNP,<br>nearest gene | Alleles (EA/OA),<br>EAF | Cancer | OR (95% CI) | P |
| --- | --- | --- | --- | --- | --- |
| 2p24.3 | rs7586503 | T/A | Breast | 0.98 (0.96-0.99) | 2.0 × 10<sup>-3</sup> |
| 15995041 | *RNU5E-7P* | 0.28 | Prostate | 0.95 (0.93-0.97) | 1.6 × 10<sup>-7</sup> |
|  |  |  | Ovarian | 0.99 (0.96-1.02) | 5.3 × 10<sup>-1</sup> |
|  |  |  | Endometrial | 0.95 (0.92-0.99) | 6.0 × 10<sup>-3</sup> |
|  |  |  | Meta-analysis<sup>c</sup> (main) |  | 6.0 × 10<sup>-9</sup> |
| 6q16.2 | rs377527229 | C/CAAA | Breast | 1.03 (1.02-1.05) | 5.9 × 10<sup>-6</sup> |
| 100091541 | *Y_RNA* | 0.23 | Prostate | 1.03 (1.01-1.05) | 1.1 × 10<sup>-3</sup> |
|  |  |  | Ovarian | 1.04 (1.01-1.08) | 6.4 × 10<sup>-3</sup> |
|  |  |  | Endometrial | 1.03 (1.00-1.07) | 8.5 × 10<sup>-2</sup> |
|  |  |  | Meta-analysis<sup>c</sup> (main) |  | 2.8 × 10<sup>-9</sup> |
| 3q27.1 | rs6784596 | G/A | ER-pos breast | 1.00 (0.98-1.01) | 7.3 × 10<sup>-1</sup> |
| 184267740 | *EIF2B5-AS1* | 0.51 | ER-neg breast | 1.05 (1.03-1.07) | 2.0 × 10<sup>-5</sup> |
|  |  |  | Prostate | 0.97 (0.95-0.98) | 8.5 × 10<sup>-5</sup> |
|  |  |  | HGSOC | 1.01 (0.98-1.04) | 5.3 × 10<sup>-1</sup> |
|  |  |  | Endometrial | 1.06 (1.03-1.09) | 2.3 × 10<sup>-4</sup> |
|  |  |  | Meta-analysis<sup>c</sup> (subtype) |  | 4.8 × 10<sup>-8</sup> |
| 17p11.2 | rs5819638 | CT/C | ER-pos breast | 1.03 (1.02-1.05) | 1.3 × 10<sup>-4</sup> |
| 17857626 | *TOM1L2* | 0.33 | ER-neg breast | 1.03 (1.00-1.05) | 2.8 × 10<sup>-2</sup> |
|  |  |  | Prostate | 1.04 (1.02-1.06) | 2.1 × 10<sup>-5</sup> |
|  |  |  | HGSOC | 1.01 (0.97-1.04) | 7.7 × 10<sup>-1</sup> |
|  |  |  | Endometrial | 1.03 (0.99-1.06) | 1.4 × 10<sup>-1</sup> |
|  |  |  | Meta-analysis<sup>c</sup> (subtype) |  | 4.9 × 10<sup>-8</sup> |

<sup>a</sup>> 1 Mb away from any previously reported lead SNP for any of the four cancers.<sup>b</sup>Build 37 position.<sup>c</sup>The Han and Eskin "RE2C\*" model does not report a combined effect size estimate and confidence interval.
Abbreviations: SNP, single nucleotide polymorphism; EA, effect allele; OA, other allele; EAF, effect allele frequency; OR, odds ratio; CI, confidence interval; ER-pos, estrogen receptor-positive breast cancer; ER-neg, estrogen receptor-negative breast cancer; HGSOC, high-grade serous ovarian cancer.

The genomic control inflation statistic scaled for 1,000 cases and 1,000 controls, λ_1000_, was 1.00 for both the main and the subtype-focused meta-analysis (**Supplementary Fig. 1**). The main meta-analysis identified 441 independent (correlated with *r*^2^ < 0.05; **Methods**) lead SNPs at genome-wide significance (*P* < 5 × 10^−8^). These spanned 183 regions at least 1 Mb apart (**Supplementary Table 2**; **Supplementary Fig. 2a**; **Methods**). The subtype-focused meta-analysis identified 393 independent genome-wide significant lead SNPs spanning 169 regions at least 1 Mb apart (**Supplementary Table 2**; **Supplementary Fig. 2b**). Two hundred and twenty-eight lead SNPs were in common to both analyses. Of the remaining 165 lead SNPs that were unique to the subtype-focused analysis, 156 SNPs were within 1 Mb of a lead SNP identified in the same region in the main analysis. Fifteen of these 156 SNPs were independent (*r*^2^ < 0.05) of the main analysis lead SNP from the same region (**Supplementary Table 2**). Nine lead SNPs were unique to the subtype-focused analysis and > 1 Mb from the main analysis lead SNPs (**Supplementary Table 2**). Thus, in total, the main and subtype-focused analyses identified 465 (441 + 15 + 9) independent lead SNPs spanning 192 (183 + 9) regions. We replicated at *P* < 5 × 10^−8^ in our current combined data set all 12 cancer susceptibility loci that we first identified in our previous multi-cancer study of breast, prostate, and ovarian cancer (**Supplementary Table 2**)^6^.

**Table 2.**
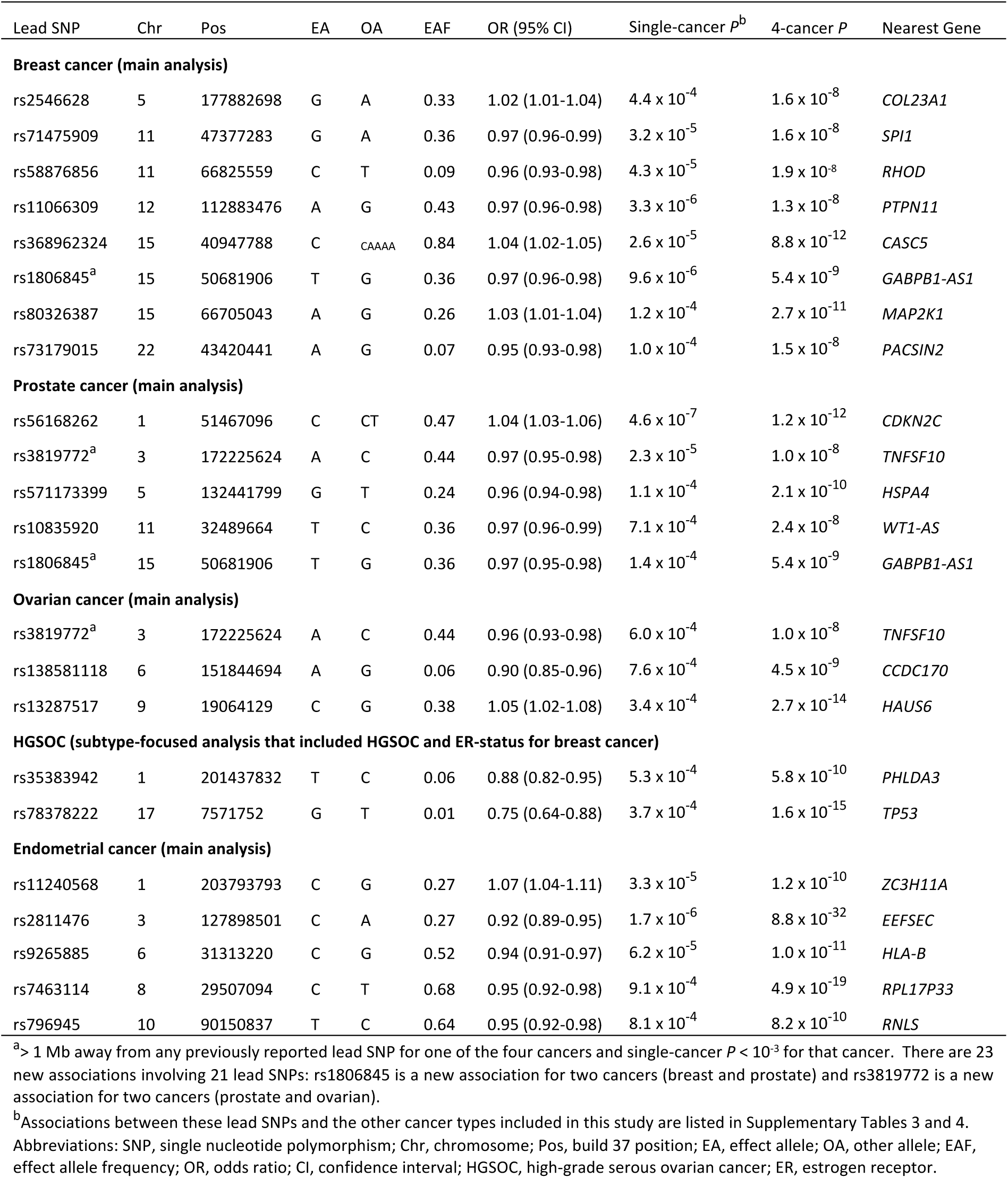
The cancer susceptibility loci not previously identified for one of the four cancers.^a^

**Fig. 1:**
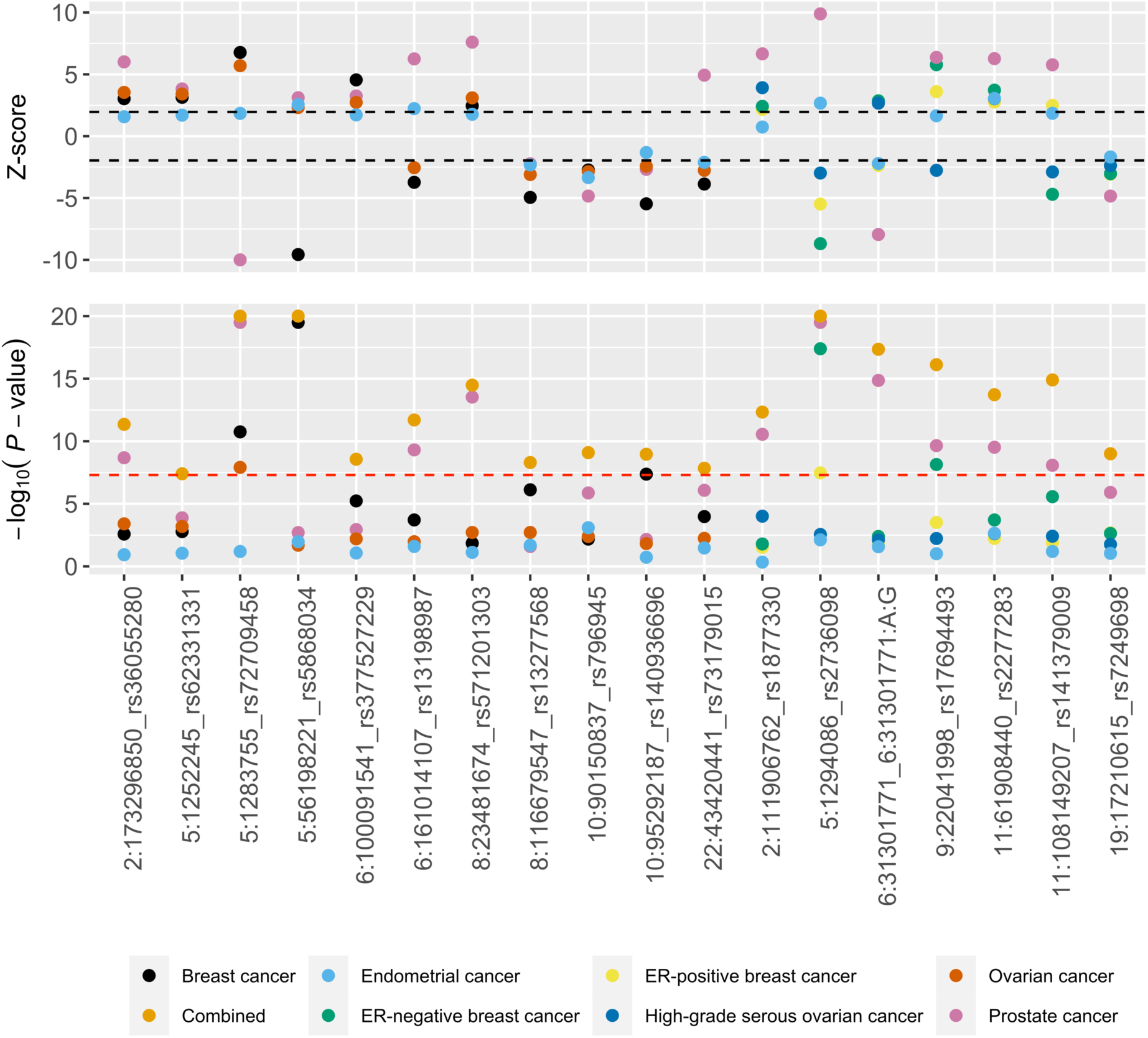
Eighteen lead SNPs in 16 regions where the Bayesian framework indicated that there was evidence for association with all four cancers out of breast, prostate, ovarian, and endometrial cancer. These lead SNPs are in 16 distinct regions at least 1 Mb apart. Eleven of the 18 SNPs were identified in the main meta-analysis and seven in the subtype-focused meta-analysis. The black dashed lines indicate Z-scores of +/- 1.96 and the red dashed line indicates a *P*-value of 5 × 10^−8^. Z-scores > 10 were set to 10 and -log10(*P*-values) > 20 were set to 20 to enable improved visualization of data points with less extreme values. “Combined” refers to -log10(*P*-values) from the Han and Eskin model-based main or subtype meta-analysis. Chromosomal positions are based on build 37.

**Fig. 2:**
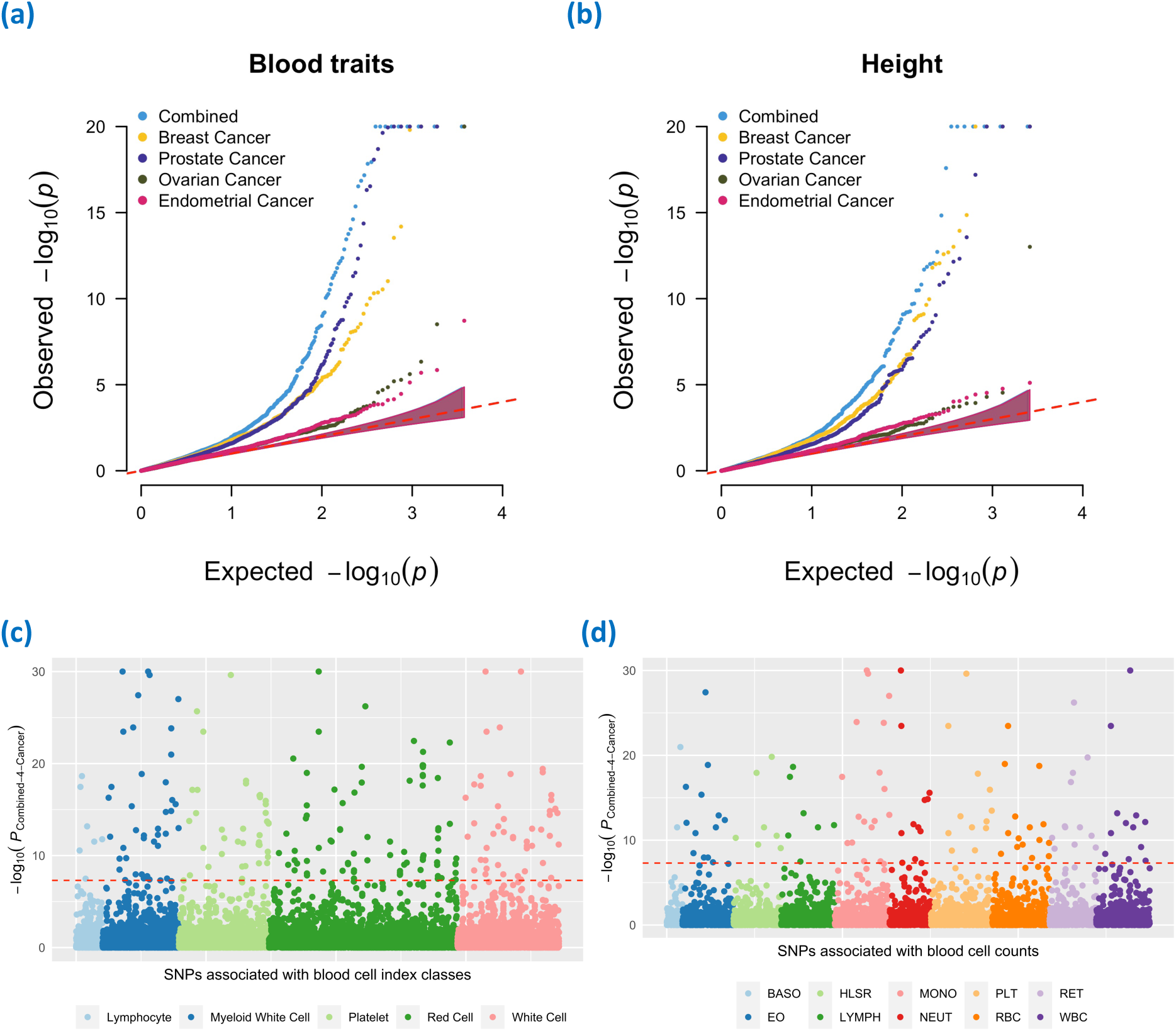
Quantile-quantile (Q-Q) plots of negative logarithm (base 10) *P*-values from the main meta-analysis (“Combined”) and the single-cancer data (breast, prostate, ovarian, and endometrial cancer) at independent variants (*r*^2^ < 0.05) associated at genome-wide significance with (a) blood cell traits in Vuckovic et al.^19^ and (b) height in Yengo et al^20^. Manhattan plot of negative logarithm (base 10) *P*-values from the main meta-analysis at variants associated at genome-wide significance in Vuckovic et al. with **(c)** blood cell index classes and **(d)** blood cell type-specific counts. Classes and counts are described in detail in Vuckovic et al. (for example, the Red Cell class includes red blood cell counts and other indices such as hemoglobin and hematocrit while the RBC count includes the red blood cell counts only). The red dashed line in the Manhattan plots indicates genome-wide significance (*P* < 5 × 10-8). Negative log10(*P*-values) > 20 were set to 20 in the Q-Q plots and > 30 were set to 30 in the Manhattan plots to enable improved visualization of data points with less extreme values. Abbreviations: BASO, basophil; HLSR, high light scatter reticulocyte; MONO, monocyte; PLT, platelet; RET, reticulocyte; EO, eosinophil; LYMPH, lymphocyte; NEUT, neutrophil; RBC, red blood cell; WBC, white blood cell.

### Cancer susceptibility loci not previously identified for any of the four cancers

We identified four lead SNPs at genome-wide significance (*P* < 5 × 10^−8^) that were at least 1 Mb away from any previously published genome-wide significant lead risk SNPs for any of the cancers evaluated in this study (**Table 1**; **Methods**). Two lead SNPs were identified from the main meta-analysis and two were identified from the subtype-focused meta-analysis.

Two additional lead SNPs, rs66686620 in the 2p13.3 region (*ANTXR1*) and rs6065253 in the 20q12 region (*MAFB*) were also genome-wide significant and over 1 Mb away from any previously reported lead risk SNP for any of the four cancers as of August 2019 (**Supplementary Table 2**). However, since then associations (*P* < 5 × 10^−8^) within this 1 Mb interval have been identified in a breast cancer genome-wide association meta-analysis by Zhang *et al*.^13^ (i) in the 2p13.3 region (rs4602255) through the addition of ∼10% more cases and ∼9% more controls to the breast cancer data set used here and (ii) in the 20q12 region (rs6065254) through the use of a statistical model that accounted for heterogeneity in associations by hormone receptor status and tumor grade in the expanded breast cancer data set. SNPs rs6065253 and rs6065254 were perfectly correlated (*r*^2^ = 1; distance between SNPs = 52bp) but rs66686620 and rs4602255 were independent (*r*^2^ = 0.003; distance between SNPs = 141kb) suggesting that the multi-cancer 2p13.3 signal is a distinct association.

### Cancer susceptibility loci not previously identified for one of the four cancers

In addition to the four loci described above, we identified 23 associations that were novel for at least one of the cancers evaluated in this study (**Table 2**). We did this by examining the lead SNPs (main and/or subtype-focused meta-analysis *P* < 5 × 10^−8^) to identify those lead SNPs that were (i) associated with one of the cancers at *P* < 10^−3^ in the contributing single-cancer data set and (ii) > 1 Mb away from any previously identified genome-wide significant lead risk SNP for the same cancer (> 10 Mb when applying this criterion to the extended major histocompatibility complex region; chr6:25— 35Mb). The 23 new associations included eight for breast cancer, five for prostate cancer, and five each for ovarian and endometrial cancer susceptibility (**Table 2**). These associations involved 21 lead SNPs since two lead SNPs, rs1806845 and rs3819772, each met these criteria for two cancer types (**Table 2**). Associations between these lead SNPs and all cancers included in this study are listed in **Supplementary Tables 3 and 4**.

**Table 3.**
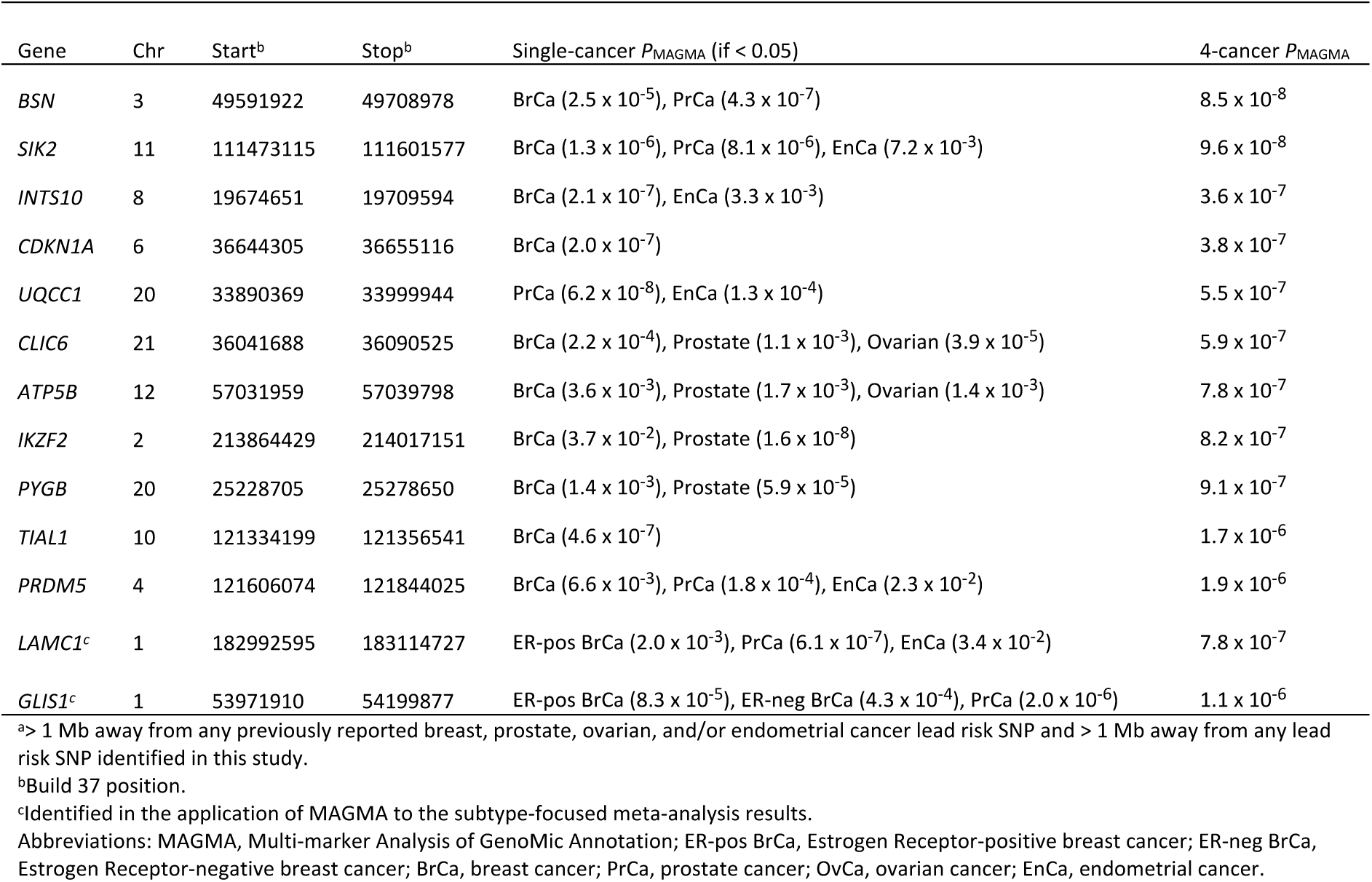
The 13 new cancer susceptibility loci identified by gene-based association (MAGMA) analysis.^a^

We identified seven associations for breast cancer (in addition to the eight reported above) that were new in the context of the breast cancer data set used in this study but have since been reported for breast cancer at *P* < 5 × 10^−8^ in the meta-analysis by Zhang *et al*.^13^ described above (five of the seven associations; **Supplementary Table 5**) or in a trans-ancestry genome-wide association meta-analysis by Shu *et al*.^15^ (two of the seven associations; **Supplementary Table 5**). Shu *et al*. combined the breast cancer data set used in this study with a data set that included over 24,000 breast cancer cases and 24,000 controls of Asian ancestry.

We also assessed the relationship between 461 lead SNPs that were within 1 Mb of one of the 463 lead SNPs previously published for breast, prostate, ovarian, and/or endometrial cancer risk and those lead SNPs (**Methods**). We found that 187 of the 461 lead SNPs were independent (*r*^2^ < 0.05) of the previously published proximal single-cancer lead SNPs (**Supplementary Table 6**). This included 8 of the 21 SNPs in **Table 2**: rs71475909, rs58876856, rs1806845, rs80326387, rs73179015, rs56168262, rs3819772, and rs9265885.

### Bayesian re-evaluation of lead variants

We used the breast, prostate, ovarian, and endometrial cancer summary genetic association data to also perform meta-analysis within a Bayesian framework^16^ for each lead SNP identified in the main and subtype-focused meta-analysis (**Methods**). This was motivated by two aims. First, to identify the subset or combination of cancers that was most likely to be responsible for the association at each lead SNP. Second, to quantify the combined evidence of association for this subset of cancers at the lead SNP in terms of the posterior probability of association (PPA), assuming a prior probability of association of 1 in 100,000. The Bayesian framework evaluated the evidence for 16 combinations at each lead SNP from the main meta-analysis (four non-pleiotropic combinations involving one cancer type only, six involving pairs and four involving triplets of cancers, one with all four cancers, and one where none of the cancers are associated) and 32 combinations at each lead SNP from the subtype-focused meta-analysis.

The Bayesian meta-analysis confirmed that the combination of cancers with greatest evidence for underpinning the association at each of the lead SNPs listed in **Table 2** included the cancer for which we report the association as novel (**Supplementary Table 7**). It also confirmed that for all but one of these lead SNPs, the PPA for this combination of cancers exceeded 80% (**Supplementary Table 7**). The only exception was rs58876856 with a PPA of 67% for the top-ranked combination that included contributions from breast, prostate, and endometrial cancer. Further, the top combination included at least three cancers for each of the four lead SNPs listed in **Table 1** and these combinations all achieved PPA > 80% (**Supplementary Table 7**). The top-ranked combination involved all the cancers included in this study for 18 lead SNPs in 16 regions (PPA > 80%; **Figure 1**). The top combination involved at least two cancers and had PPA > 80% for 222 of the 465 lead SNPs (in 118 of the 192 regions; **Supplementary Table 8** and **Methods**). The associations at the remaining lead SNPs were driven by a single cancer or the combination of ER-positive and ER-negative breast cancer (**Supplementary Table 8**).

### Associations between lead variants and other diseases and traits

We investigated whether the 465 lead SNPs identified in this study or SNPs in strong linkage disequilibrium (LD; *r*^2^ > 0.8) with them were associated with other diseases and traits at genome-wide significance (*P* < 5 × 10^−8^) in previously published studies (**Methods**)^17^. We found that 97/465 lead SNPs (or SNPs in strong LD) were associated with at least one trait or disease (other than breast, prostate, ovarian, and/or endometrial cancer; **Supplementary Table 9**). These associated traits or diseases were classified in terms of broader categories (specifically, Experimental Factor Ontologies or EFOs)^18^. The 97 SNPs were associated with 190 unique EFOs (**Supplementary Table 9**). The two EFOs associated with the largest number of lead SNPs were EFO_0004586 (“complete blood cell count”; 54/97 SNPs, including 42/222 SNPs that were associated with at least two cancers based on the Bayesian evaluation) and EFO_0004339 (“body height”; 10/97 SNPs). A recent (unpublished) study^19^ has substantially increased the number of genomic loci associated with blood cell phenotypes and there are now 16,900 known genetic associations involving 29 blood cell traits. Having shown that ∼12% of the lead SNPs from our study (or nearly a fifth of the lead SNPs with detectable cross-cancer effects in the Bayesian evaluation) were associated with blood cell traits (EFO_0004586), we asked if the converse was true: were genome-wide significant SNPs from the recent blood cell trait GWAS also enriched for low *P*-values in the main four-cancer meta-analysis? We found that there was evidence for such enrichment (**Fig. 2a**), the enrichment was driven by all four cancers but particularly breast and prostate cancer (**Fig. 2a**), and the enrichment closely paralleled that seen at variants known to be associated with height (**Fig. 2b**)^20^, another highly polygenic quantitative trait that ranked second in our EFO list. We found that this enrichment was not restricted to SNPs associated with a particular blood cell index class as classified by Vuckovic et al.^19^ (**Fig. 2c**) or with the counts of a particular blood cell type (**Fig. 2d**).

### Functional annotation of the new lead SNP-cancer associations

We annotated the lead SNPs (and SNPs in strong LD; *r*^2^ > 0.8) marking the four loci not previously identified for any of the four cancers and the 23 new lead SNP-cancer associations using several genomics resources to identify gene targets with functional support (**Methods**)^21–28^. This included expression quantitative trait loci (eQTLs) identified at false discovery rate (FDR) < 0.05 by the Genotype-Tissue Expression (GTEx) consortium^22^ in breast, prostate, ovary, and uterus (*cis*-eQTLs; **Supplementary Table 10**) and by the eQTLGen consortium^23^ in blood (*cis*- and *trans*-eQTLs; **Supplementary Table 11**); and Combined Annotation Dependent Depletion (CADD) scores^24^ (**Supplementary Table 12**). Among the protein coding genes nearest to the 27 lead SNPs, the GTEx database supported *COL23A1, HLA-B, HAUS6, RHOD*, and *GABPB1* as targets (**Supplementary Table 10**) while the eQTLGen database supported *COL23A1, HAUS6, RHOD, EPHB3, CCDC170, EEFSEC, HSPA4, MAP2K1, PACSIN2, RNLS, SPI1, TP53*, and *ZC3H11A* as targets (**Supplementary Table 11**). Lead SNP rs35383942 in an exon of *PHLDA3* had a CADD score of 24. This SNP is a missense variant (p.Arg28Gln) known to be associated with increased breast cancer risk^7^ that we found associates with decreased ovarian cancer risk (OR = 0.88, 95% CI: 0.82—0.95; *P*_HGSOC_ = 5.3 × 10^−4^).

### Direction of allelic association across pairs of cancers

Next, we examined the 441 genome-wide significant lead SNPs from the main analysis and the 393 genome-wide significant lead SNPs from the subtype-focused analysis to characterize the direction of allelic association across cancers with an emphasis on SNPs that were strongly associated with at least two cancers individually (*P* < 10^−3^). We considered every possible pairwise combination of cancers in the main and subtype-focused analyses (except the pairing of ER-positive breast cancer with ER-negative breast cancer). We observed that for 94 of the 441 main lead SNPs (**Supplementary Table 3**) and 89 of the 393 subtype-focused lead SNPs (**Supplementary Table 4**), the lead SNP was associated with each cancer out of at least two cancers at *P* < 10^−3^ in the corresponding single-cancer data set. We focused further analyses presented in this section of the results on these subsets of lead SNPs. For 29/94 main lead SNPs and 33/89 subtype lead SNPs, the allele that conferred risk of developing one of the two cancers had a protective association with the other cancer (**Supplementary Table 13**).

We explored the biological basis of these alleles that have opposite effects across cancers by analyzing over-representation^29^ of pathways from the Kyoto Encyclopedia of Genes and Genomes (KEGG)^30^ among the combined set of genes nearest to the lead SNPs that showed opposite effects across cancer pairs in the main and subtype-focused analysis. The strongest enrichment observed was for genes in the KEGG p53 signaling pathway (FDR = 4.4 × 10^−5^; **Supplementary Table 14**). This p53 pathway enrichment was underpinned by five genes: *ATM, CASP8, CHEK2, MDM4*, and *TP53*. We visualized single-cancer associations at lead SNPs/alleles in or nearest to these five genes using a forest plot (**Fig. 3**). In contrast, we observed a distinct pattern of pathway over-representation (**Supplementary Table 14**) among the combined set of genes nearest to the lead SNPs that showed same direction associations (single-cancer *P* < 10^−3^) across cancer pairs in the main analysis (65/94 lead SNPs; **Supplementary Tables 3 and 13**) and subtype-focused analysis (56/89 lead SNPs; **Supplementary Tables 4 and 13**). The KEGG apoptosis pathway was the highest-ranked pathway shared between the opposite and same direction analyses (**Supplementary Table 14**).

**Fig. 3:**
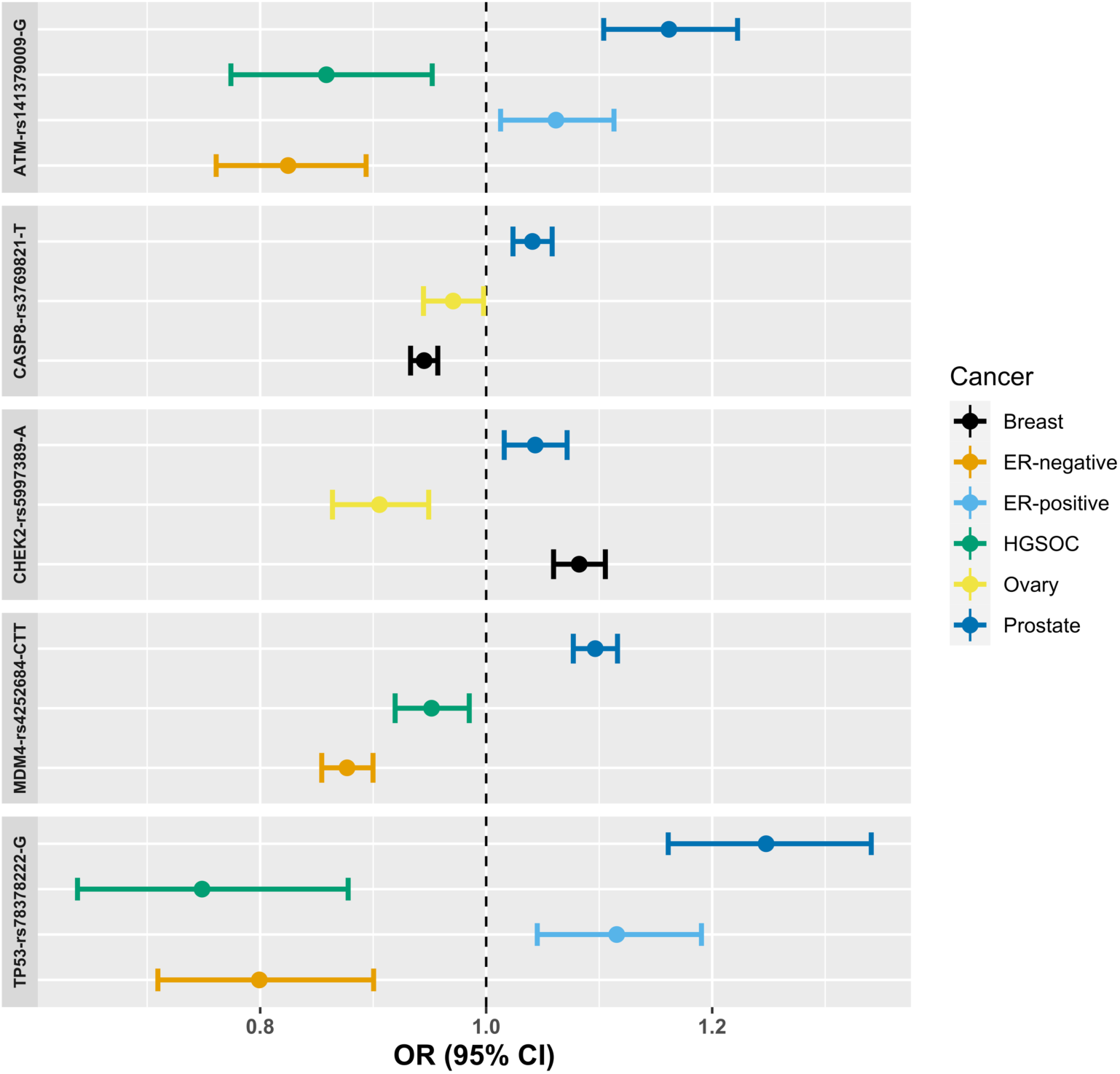
Forest plots showing odds ratios (OR) and 95% confidence intervals (CI) for the lead SNPs/effect alleles in or nearest to the five genes in the TP53 signalling pathway that underlie the enrichment for this pathway. All cancer- or subtype-specific associations at *P* < 0.05 are shown in the forest plot. Abbreviations: ER, estrogen receptor; HGSOC, high-grade serous ovarian cancer.

### Gene-level association analysis

We mapped all SNPs along with their association *P*-values from the main and subtype-focused genome-wide association meta-analyses to 19,100 protein-coding genes across the genome and performed gene-level association testing using Multi-marker Analysis of Genomic Annotation (MAGMA^31^). MAGMA collapses multiple SNP-level associations in a gene while taking into account LD between the SNPs and offers a gene-level association test that complements conventional single-SNP association analyses.

Before proceeding further, we evaluated the biological meaningfulness of the gene-level MAGMA associations using gene expression levels in 54 tissue types from the GTEx database^22^. Expression levels of genes in the breast, prostate, fallopian tube, and uterine tissue are predictors of the gene-level multi-cancer MAGMA association signals (*P* < 1.7 × 10^−5^; **Supplementary Tables 15 and 16**). This finding is consistent with these tissues being relevant to the etiology of the multi-cancer phenotype. A much weaker association was observed between gene expression in ovarian tissue and the gene-level MAGMA signal (*P* = 0.03). We also applied MAGMA to the breast, prostate, ovarian, and endometrial cancer genome-wide SNP association data sets individually. The corresponding single cancer gene-level signals were less strongly associated with tissue-specific gene expression levels than the multi-cancer gene-level signals for breast (*P*_single_ = 1.7 × 10^−5^ and *P*_multi_ = 1.5 × 10^−7^), fallopian tube (*P*_single_ = 0.01 and *P*_multi_ = 1.3 × 10^−6^), ovary (*P*_single_ = 0.06 and *P*_multi_ = 0.04), and uterus (*P*_single_ = 0.03 and *P*_multi_ = 2.6 × 10^−6^), but not in the prostate (*P*_single_ = 3.9 × 10^−18^ and *P*_multi_ = 5.6 × 10^−15^). This indicates that the gain in power obtained by combining data across cancers was, in general, improving the association signal for genes more likely to be expressed in the relevant tissue types compared to single cancer analyses and suggests the presence of shared mechanisms underlying inherited cross-cancer risk acting across multiple anatomically separate tissue sites.

The multi-cancer gene-based association analysis identified 13 genes in as many regions (**Table 3**) at genome-wide significance (*P*_MAGMA_ < 2.62 × 10^−6^ after accounting for testing 19,100 genes) that were > 1 Mb away from any previously identified breast, prostate, ovarian, and/or endometrial cancer lead risk SNP and > 1 Mb away from any lead SNP identified in SNP-based analysis in this study.

### Pathway analysis based on gene-level associations

We performed pathway analysis on the set of genes with gene-level association *P*_MAGMA_ < 2.6 × 10^−6^ from the main meta-analysis or from the subtype-focussed meta-analysis and in the top 5% of the ranked gene-level associations for at least two cancers from the single-cancer analyses. Three hundred and thirteen genes met these criteria from which we excluded five HLA genes (**Supplementary Tables 17**). The final set of 308 genes was enriched for twelve pathways from KEGG at FDR < 0.05 with over five-fold enrichment in two pathways: p53 signalling and endocrine resistance (**Supplementary Tables 18**). The enrichment in the p53 signalling pathway was driven by *CCND1, CDKN2A, SIVA1, CCNE1, CASP8, CHEK2*, and *MDM4* and was more modest than the enrichment for the same pathway observed in the 37 genes nearest to the set of SNPs with opposite effects on at least two cancers, which was driven by *TP53, ATM, CASP8, CHEK2*, and *MDM4*. Performing the pathway analysis retaining the five *HLA* genes (i.e., using all 313 genes) implicated several immune-related pathways where the signal was largely driven by these genes (**Supplementary Tables 19**).

## DISCUSSION

We performed the largest genome-wide association meta-analysis for shared susceptibility to four hormone-related cancers – breast, prostate, ovarian, and endometrial cancer – to date. This identified four new risk loci that were over a megabase away from previously identified risk loci for the four cancers and 21 new risk loci incorporating 23 new lead SNP-cancer type associations that were over a megabase away from previously identified risk loci for at least one of the cancers. Gene-based association analysis identified an additional 13 novel risk loci. Beyond locus discovery, we found evidence for pleiotropic cross-cancer effects at ∼48% (222/465) of the genome-wide significant lead SNPs in our study. The strongest association model at these SNPs included contributions from at least two of the four cancers. We identified several risk loci where the lead SNP had strong associations with at least two of the cancers but opposite effects (for the same effect allele) on these cancers. We demonstrated enrichment of the multi-cancer association signal in breast, prostate, fallopian tube, and uterine tissues, conducted a phenome-wide association scan of cross-cancer lead SNPs revealing profound overlap with blood cell trait loci, and observed enrichment of pivotal pan-cancer pathways in our combined data set.

The 8q24.1 region, a “gene desert” with a paucity of protein coding genes other than *MYC* was one of the first regions to be recognized as one that contains SNPs associated with multiple cancer types^32^. Here we report a novel cancer susceptibility locus in a similar gene desert, 2p24.3 with lead SNP rs7586503 and nearest coding gene *MYCN*, which shows evidence for association with breast, prostate, and endometrial cancers (**Table 1**). The proto-oncogene *MYCN* is known for its ability to drive the development of multiple tumor types that arise from a common cellular lineage^33^. The lead SNP rs5819638 in the 17p11.2 region is an intronic insertion and deletion variant in *TOM1L2* (**Table 1**). This SNP is correlated (*r*^2^ = 0.62) with rs8070624, a known genome-wide significant lead SNP for bone mineral density (BMD)^34^, an estrogen-regulated trait in both men and women^35^. The T allele deletion of rs5819638 is protective for breast, prostate, and endometrial cancer. The deletion corresponds to the G allele of rs8070624 that is associated with reduced BMD, which is consistent with the observed relationship between lower BMD and decreased breast, prostate, and endometrial cancer risks^36–38^. The A allele of the new breast cancer risk lead SNP rs71475909 increases breast cancer risk but decreases prostate cancer risk (**Table 2** and **Supplementary Table 3**). The SNP lies in an intron of *SPI1*, which encodes a transcription factor that co-binds with TP53 to TP53 target genes and is crucial for the maintenance of pro-apoptotic transcriptional repression by TP53^39,40^. Expression QTL analyses in breast, prostate, ovarian, uterine, and whole blood tissue types supported several genes as candidate targets at the new risk loci. Lead SNP rs58876856 was associated with breast (*P* = 4.3 × 10^−5^; **Table 2**) and prostate (*P* = 4.9 × 10^−4^; **Supplementary Table 3**) cancer risk and with *RHOD* expression in breast (*P* = 6.4 × 10^−25^) and prostate (*P* = 1.3 × 10^−16^) tissues. *RHOD* encodes a member of the Rac subfamily of the Rho family of GTPases that has an established oncogenic role^41^. We also identified a novel association for HGSOC risk at rs35383942, a known genome-wide significant lead SNP for breast cancer risk^7^ and male-pattern baldness^42^. The T allele of rs35383942 is associated with increased susceptibility to breast cancer but decreased susceptibility to HGSOC. SNP rs35383942 is a missense variant (CADD score 24; **Supplementary Table 12**) in *PHLDA3*, a tumor supressor gene that sits at the intersection of two key cancer pathways: PHLDA3 is directly regulated by TP53 and is a repressor of AKT1 signaling, contributing to TP53-dependent apoptosis^43^. The new endometrial cancer risk SNP rs2811476 (**Table 2**; at a known prostate cancer risk locus – **Supplementary Table 3**), in an intron of *EEFSEC*, was strongly correlated with rs2955117 (*r*^2^ = 0.95) and rs2687729 (*r*^2^ = 0.99), which are genome-wide significant lead SNPs for the hormone-related traits maternal effect on gestational age at birth^44^ and age at menarche^45^, respectively. The alleles associated with longer gestational duration and later menarche were protective for endometrial cancer and these findings are consistent with the “unopposed oestrogen hypothesis” for endometrial cancer^46^. The gene-based MAGMA approach identified 13 new candidate cancer susceptibility genes in 13 loci not detected by SNP-level single- and cross-cancer genome-wide association meta-analyses. Each of these genes is discussed in detail in **Supplementary Note 1**.

We identified 18 lead SNPs in 16 regions where a Bayesian evaluation of the data suggested that the combination of cancers with greatest evidence for association involved all four cancers. Some of these lead SNPs marked known multi-cancer susceptibility loci near *TERT*^47^, CDKN2B-AS1^48^, *INCENP*^6^, and *BCL2L11*^6^. Other regions (out of the 16) have not been recognized before for such extensive cross-cancer pleiotropic effects. For example, lead SNP rs5868034 is located ∼7kb from *SETD9* which encodes a lysine methyltransferase that methylates TP53, increasing its stability and promoting the expression of its targets *CDKN1A, BAX*, and *MDM2*^49^. While rs5868034 is ∼144kb from rs62355902, a known breast cancer risk lead SNP that regulates *MAP3K1* in the same region^50^, the two SNPs are independent (*r*^2^ = 0.03). Further, a GTEx look-up for rs5868034 shows that it is associated with the expression of *SETD9* in breast (*P* = 1.9 x 10^−32^), prostate (*P* = 8.5 × 10^−18^), ovarian (*P* = 9.6 × 10^−12^), and uterine (*P* = 1.6 × 10^−11^) tissues^22^. Lead SNP rs140936696 is ∼3kb from *CEP55*, which encodes a centrosome- and midbody-associated protein that is a major regulator of abscission or the final stage of cytokinesis^51^ and of the PI3K/AKT pathway^52^. *CEP55* is a member of several common gene expression signatures (such as the “CIN70”) that serve as indices of chromosomal instability, cell cycle progression, proliferation, and metastatic potential across multiple cancer types^53^. Lead SNP rs796945 near *RNLS* is associated with breast, prostate, ovarian, and endometrial cancers and has been recently identified as a lead SNP for the hormone-related gynecologic disorder endometriosis^54^, all with the same direction of allelic effect.

We observed a substantial overlap between the cross-cancer genetic associations from this study and variants known to be associated with a range of blood cell phenotypes. Some loci associated with blood cell traits, in particular blood cell counts, likely represent cell-type agnostic biomarkers of cellular turnover and proliferative potential – biological processes that, in turn, may link them to solid tumor risk. Indeed, common genetic variants at *SH2B3, ATM, TERT, TET2*, and *CHEK2* are associated with multiple blood cell traits^55^ and with myeloproliferative neoplasms^56,57^, and were the genes nearest to five cross-cancer lead SNPs identified in the current study. For example, our lead SNP rs7310615 that maps to *SH2B3* (*P*_breast_ = 3 × 10^−7^ and *P*_endometrial_ = 1.3 × 10^−10^) is a genome-wide significant lead SNP associated with white blood cell, red blood cell and platelet counts and other blood cell traits^55^, and with myeloproliferative neoplasms^56,57^. Rare germline missense variants in *SH2B3* and *ATM* are also associated with the blood cell (somatic) acquisition of loss-of-heterozygosity (LOH) in the chromosomal arms where these genes reside, which may result in the aberrant clonal expansion of the blood cells acquiring these LOH events or clonal hematopoeisis^58^.

Genes nearest to cross-cancer risk lead SNPs in this study – *BCL2L11, SMC2, CTSK, USP28, CDKN1B, MAP2K1, TP53, ATM*, and *CHEK2* – also mark loci predisposing to mosaic Y chromosome loss in blood, the most common form of clonal hematopoeisis^59^. Overall, these genetic overlaps with blood cell traits add to a growing body of evidence which suggests that clonal hematopoiesis may be a proxy for genomic instability in the body that is associated with non-hematological cancer risk^59^.

Pathway analysis of our 2016 breast-prostate-ovarian cancer shared susceptibility GWAS meta-analysis findings had identified a central role for induction of apoptosis through death receptor signaling^6^. In our current substantially expanded data set that also includes endometrial cancer we not only continue to detect a signal for the apoptosis pathway but also highlight roles for endocrine resistance and TP53 signaling. We specifically note a pattern involving cross-cancer susceptibility alleles at the TP53 signaling genes *CASP8, MDM4, CHEK2, ATM* and *TP53* wherein, in general, lead SNP alleles conferring risk of ovarian cancer/HGSOC and ER-negative breast cancer show protective associations with overall/ER-positive breast cancer and prostate cancer. This pattern likely reflects the differential prevalence of loss of tumor suppressor function versus dominant negative or gain-of-function (GOF) oncogenic *TP53* somatic mutations that is seen in these tumor types^60,61^. For example, the G allele of lead SNP rs78378222 is associated with lower expression of *TP53* (*P* = 6 × 10^−21^ in eQTLGen^23^; **Supplementary Table 11**). This SNP is known to alter the polyadenylation signal and impair the 3′-end processing of *TP53* mRNA^62^. The G allele is protective for HGSOC and ER-negative breast cancer (**Fig. 3**), where dominant negative or GOF *TP53* somatic mutations are more common, potentially because the lower germline-regulated expression attenuates the oncogenic effects of TP53 when such a mutation is somatically acquired. However, the same allele confers susceptibility to ER-positive breast and prostate cancer (**Fig. 3**), where loss-of-function (LOF) *TP53* somatic mutations predominate, potentially because the lower background expression accentuates the loss of TP53 tumor suppressor function when the LOF mutation is somatically acquired. Thus, our findings hint at a potential germline-somatic interaction that strongly warrants further investigation. Antagonistic effects across cancer types at the same allele also has important implications for the use of such alleles in the development and application of polygenic risk scores that have so far been confined to a consideration of allelic associations with a single cancer. The identification of alleles with opposite effects across the cancers was helped by that fact that the Han and Eskin model^11,12^ used in this study can detect effects under heterogeneity in contrast to the standard fixed-effects model used in the 2016 meta-analysis^6^. Opposite associations across related diseases at the same allele are also seen in autoimmune^63^ and psychiatric^64^ disease groups. While a direct comparison between cancer, autoimmune, and psychiatric studies is challenging due to differences in sample size (the cancer data set here is larger by case numbers) and design, the prevalence of such associations in cancer appears to be higher.

It is worth noting that 187 of the 465 lead SNPs highlighted in this study were within 1 Mb of known breast, prostate, ovarian, and/or endometrial cancer risk loci but not linked (*r*^2^ < 0.05) to the published lead SNPs at these loci. This finding is consistent with a recent combined GWAS of multiple blood lipid traits^65^ that identified several independent (in terms of LD) associations at known lipid-associated loci. These associations were not reported in the corresponding single lipid GWAS. This suggests that for related traits (groups of cancers or groups of lipids), the multi-trait genetic architecture may, at least in part, be distinct from the single-trait genetic architecture. That is, the same genomic regions may harbor distinct SNPs that have disease-specific and cross-disorder associations. This is best illustrated with an example: rs58058861 is a known genome-wide significant lead SNP for breast cancer risk^7^ and our meta-analysis identifies an independent (*r*^2^ = 0.01; distance between SNPs = 60kb) lead SNP rs3819772 in the same region and ∼16kb from *TNFSF10* that is associated with breast cancer risk albeit less strongly (*P* = 2.8 × 10^−4^: **Supplementary Table 3**). However, rs3819772 from our meta-analysis is pleiotropic (**Table 2**) and also associates with prostate (*P* = 2.3 × 10^−5^) and ovarian (*P* = 6 × 10^−4^) cancer risk. *TNFSF10* encodes TRAIL, a cytokine of the tumor necrosis factor ligand superfamily that is expressed in most normal tissues and selectively induces apoptosis in transformed and tumor cells by binding to death receptors^66^. Subsequent studies should aim to validate the complex multi-trait genetic architecture in such regions by orthogonal approaches such as evaluating SNP interactions with tissue-specific and cross-tissue gene regulatory mechanisms.

In our investigation of the relationship between gene-level associations and tissue-specific gene expression we found that tissue-specific gene expression in the ovary and fallopian tube are similarly and weakly associated with gene-level associations based solely on the ovarian cancer data set used here. However, gene-level associations derived from the four-cancer meta-analysis are much more strongly associated with tissue-specific gene expression in the fallopian tube than in the ovary. This result is consistent with emerging evidence that the cell of origin of HGSOCs, the most common and aggressive histotype of ovarian cancer, is in the fallopian tube^67,68^, and suggests that combining data from other cancers improves the ranking of genes associated with ovarian cancer risk that are also fallopian tube-specific. We identified nine lead SNP-breast cancer associations in our study that were new in the context of the breast cancer data set used here but have recently been identified in two larger breast cancer GWAS^13,15^. These breast and ovarian cancer findings, in particular, demonstrate that cross-cancer GWAS meta-analysis can be a powerful approach to the identification of new cancer susceptibility loci. The approach can be comparable, for pleiotropic regions, to the addition of cases and controls from the same cancer type and boost the discovery of gene-level signals that are expressed in the most relevant tissues. This is of particular relevance for elucidating the genetic architecture of rarer cancer types where larger GWAS studies may not be feasible.

In summary, this large-scale meta-analysis combining GWAS data for breast, prostate, ovarian, and endometrial cancers identified four shared susceptibility loci not previously reported for any of the four cancer types and 23 lead SNP-cancer associations in 21 shared susceptibility loci that were new for at least one of the four cancer types. Examining these data under a gene-level lens demonstrated that multi-cancer gene-level associations were strongly enriched in the relevant tissues and yielded an additional 13 loci containing candidate breast, prostate, ovarian, and/or endometrial cancer susceptibility genes. Bayesian analysis generated the first comprehensive genome-wide catalog of lead SNPs where the association was likely to be driven by at least two of these cancers. We obtained fresh insights into the biology of the shared susceptibility loci, particularly those alleles that displayed opposite associations across cancers. Potential target genes and other phenotypic associations at the shared susceptibility loci suggest a complex interplay between inherited hormone-related cancer risk and major molecular mechanisms acting across cancer sites. These cross-cancer risk loci provide a rich substrate for the future development of laboratory-based functional studies imperative for the translation of discoveries from this GWAS into clinical and public health impact across cancer types.

## METHODS

### GWAS data, data harmonization, and meta-analysis

SNP genotype and sample quality control, ancestry inference, imputation, genome-wide association and meta-analysis procedures for the breast, prostate, ovarian, and endometrial cancer GWAS meta-analysis data sets have been described previously^7–10^. All analyses were based on individuals of European ancestry and used 1000 Genomes Phase 3 (Version 5)-imputed or genotyped SNPs^69^. We harmonized effect and non-effect alleles and effect size estimate (beta coefficient) signs across the GWAS meta-analysis summary statistics data sets included in this study so that beta coefficients were signed based on the same effect allele in all data sets. We focused all analyses on 9,530,997 variants with minor allele frequency > 1% and imputation quality > 0.4 in the largest data set (breast cancer).

Meta-analysis was carried out using the Han and Eskin^11^ “RE2”model that has been shown to offer greater power to detect associations with heterogenous effects across traits in the context of GWAS as compared to the conventional DerSimonian and Laird^70^ random-effects model which was originally developed for the meta-analysis of clinical trials. The “2” in “RE2” refers to the fact that the RE2 test statistic has two parts, a part to detect heterogeneity (random effects) and a mean effect part that is equal to the square of the fixed-effects meta-analysis test statistic^11^. The latter enables RE2 to detect associations where the effect size estimates are homogeneous across studies. We specifically used a recently developed extension of the RE2 model that was able to take into account correlation between GWAS summary statistics due to overlapping controls. This extension is referred to as “RE2C*” and was implemented using the RE2C version 1.04 software^12^.

Correlation between GWAS summary statistics due to overlapping controls was estimated using the tetrachoric correlation between binary-transformed GWAS summary z-scores (z ≥ 0 classified as 1 and z < 0 classified as 0)^71,72^. The tetrachoric correlation was computed using the Digby approximation (implemented in Stata 14.0) as recommended by Southam et al^71^. Southam et al. have previously demonstrated the accuracy of the tetrachoric correlation for estimating the correlation between GWAS summary statistics due to overlapping controls^71^. RE2C* uses the correlation between GWAS summary statistics due to overlapping controls to inflate the standard errors of, and “decouple”^73^, the GWAS summary statistics included in a meta-analysis. Thus, the effective sample size of the RE2C* meta-analysis is the sample size obtained by simply adding up all the contributing GWAS data set sample sizes. The RE2C* model does not report a combined effect size estimate and confidence interval and only provides a *P*-value.

### Independent lead SNPs, genomic regions, and known cancer susceptibility loci

The random-effects meta-analysis results were processed using the FUMA integrative platform^21^. To define independent lead SNPs, the relevant FUMA SNP2GENE tool parameters were set as (1) Minimum P-value of lead SNPs < 5 × 10^−8^, (2) *r*^2^ threshold to define lead SNPs ≥ 0.05, (3) Reference panel population of 1000G Phase3 EUR, and (4) maximum distance between independent lead SNPs to merge into a region set as < 1000kb (i.e., lead SNPs closer than this distance were merged into a single genomic region). We also used FUMA SNP2GENE to assess the relationship between 463 lead SNPs previously reported for breast, prostate, ovarian, and/or endometrial cancer risk and the 461 lead SNPs within 1 Mb of these SNPs identified in the main and subtype-focused analyses (same parameters used as above except the maximum distance parameter that was set to 0kb). Only variants that had either been genotyped or imputed with a quality score > 0.8 were used to define lead SNPs. A comprehensive list of previously reported cancer susceptibility loci for breast, prostate, ovarian, and endometrial cancers (including susceptibility loci known to be shared by at least two of these cancers) was drawn up based on lead SNPs reported in 11 publications^6–10,13,15,48,74–76^.

### Bayesian meta-analysis

The exhaustive subset search function in the MetABF R package^16^ was used to perform the Bayesian re-evaluation of the combined association at each lead SNP identified in the main- and subtype-focused Han and Eskin RE2C* model-based meta-analyses. The exhaustive subset search function was used to evaluate the evidence for all 16 possible combinations at each lead SNP from the main meta-analysis (four non-pleiotropic combinations involving one cancer type only, six involving pairs and four involving triplets of cancers, one with all four cancers, and one where none of the cancers are associated) and, similarly, all 32 possible combinations at each lead SNP from the subtype-focused meta-analysis. MetABF, like RE2C*, takes into account the correlation in GWAS summary statistics due to overlapping controls. This is referred to as “cryptic correlation” in the MetABF tool and the same tetrachoric correlation matrix applied to the RE2C* meta-analysis was used for the MetABF meta-analysis. Further, MetABF allows the specification of a prior expected correlation. This was set to 0 to parallel the ability of RE2C* to detect heterogeneous or random effects and set to 1 to parallel the ability of RE2C* to detect homogeneous or fixed effects and the results were averaged for the two prior correlations specified. MetABF also allows for the specification of a prior expected upper bound on the effect size estimate (odds ratio) for each study (in this case, each cancer type-specific GWAS). For example, if this prior odds ratio is set to 1.5, it reflects a belief that the prior probability that the odds ratio is larger than 1.5 is 2.5%. We used each cancer-specific summary genetic association data set to determine this prior by filtering the data set to retain only the SNPs achieving *P* < 5 x 10^−8^ and calculating the 97.5^th^ percentile of the odds ratios. Thus, this reflected the odds ratios for SNPs associated with each cancer detectable given the sample size available for the cancer. The approximate Bayes factors (ABFs) calculated by MetABF were converted into posterior probabilities of association (PPAs) using the formula, PPA = (ABF × 10^−5^)/(ABF × 10^−5^ + 1), assuming a prior probability of association at each SNP of 1 in 100,000 or 10^−5^.

### Associations between lead SNPs and other diseases and traits

The PhenoScanner version 2 bulk search tool (search date 29 May 2020) was used with query set to SNP, catalogue to diseases and traits, p-value to 5E-8, proxies to EUR, *r*^2^ to 0.8, and build to 37. Only associations reported in publications with a PubMed identifier (PMID) were retained in the output.

Lists of genetic associations with blood cell traits and height were obtained: the Yengo et al.^20^ Genetic Investigation of Anthropometric Traits (GIANT) consortium file “Meta-analysis Wood et al + UKBiobank 2018 top 3290 Height SNPs from COJO analysis GZIP” was downloaded (link in the URLs section) for height and *Supplementary Table 3* from Vuckovic et al.^19^ was downloaded for blood cell traits. The FUMA pipeline (with settings as described above) was used to define independent (in terms of LD) SNPs from these genetic association files. Association *P*-values from the main four cancer and each of the cancer-specific meta-analyses at the resultant SNP positions and the CMplot R package (version 3.6.0) were used to generate the quantile-quantile plots in **Fig. 2**. Blood cell trait-associated SNP position and blood cell index class and count information from *Supplementary Table 3* from Vuckovic et al.^19^ and association *P*-values from the main four-cancer meta-analysis were used to generate the Manhattan plots in **Fig. 2**.

### Pathway analyses, functional and nearest gene annotation

All pathway analyses were conducted using the over-representation analysis function in WebGestalt (web-based gene set analysis toolkit)^29^ and pathways from the Kyoto Encyclopedia of Genes and Genomes release 88.2, 11/01/2018^30^. Functional annotation and nearest gene identification were also undertaken using the FUMA pipeline and included expression quantitative trait loci (eQTLs) identified at false discovery rate (FDR) < 0.05 by the Genotype-Tissue Expression (GTEx) consortium (version 8 data set; *cis*-eQTLs)^22^ in breast (n = 396), prostate (n = 221), ovary (n = 167), and uterus (n = 129; uterine but not specifically endometrial tissue) and by the eQTLGen consortium^23^ in blood (*cis*- and *trans*-eQTLs; n = 31,684), Combined Annotation Dependent Depletion (CADD) scores^24^, RegulomeDB scores^25^, ANNOVAR categories^26^, and ChromHMM states^27,28^. LDlink was used for all *r*^2^ calculations presented in the Discussion section and these were based on 1000 Genomes Project Phase 3 (Version 5) European ancestry populations^77^.

### Gene-level association analysis and tissue-specific enrichment

Gene-level association analysis was performed using the MAGMA (multi-marker analysis of genomic annotation)^31^ tool implemented in FUMA that takes into account multiple SNP-level summary genetic associations and LD between SNPs. SNPs were mapped to a gene if they were located between the start and end sites of the gene based on build 37 SNP and gene positions.

MAGMA “gene-property analysis” was performed to test for associations between gene-level signals and tissue-specific gene expression profiles. Gene-property analysis uses a multi-variable regression model that includes gene expression in a specific tissue type and the average gene expression across 54 tissue types to evaluate the relationship between tissue specificity and gene-level association. Tissue specific gene expression data were from GTEx version 8. GTEx eQTL results are only available for tissues with > 70 matched germline genotype-gene expression samples and the fallopian tube tissue samples in GTEx did not reach this sample size threshold. Therefore, fallopian tube was not included in the eQTL annotation but was available for the MAGMA gene-property analysis.

## URLs

Links to download data sets and analytic tools used in the study:

All genome-wide summary association statistics linked to this paper (input single-cancer data, matrices, priors, and output multi-cancer data): https://doi.org/10.5281/zenodo.3911767

RE2C: http://software.buhmhan.com/RE2C/index.php

FUMA and MAGMA: https://fuma.ctglab.nl/

MetABF: https://github.com/trochet/metabf

PhenoScanner: http://www.phenoscanner.medschl.cam.ac.uk/

GIANT consortium data: https://portals.broadinstitute.org/collaboration/giant/index.php/GIANT_consortium_data_files

WebGestalt: http://www.webgestalt.org/

## CONTRIBUTIONS

Study conception and design, statistical analysis, data interpretation, writing of first draft, guarantor: S.P.K.; Study conception, data interpretation, writing: S.L., R.J.Hu., P.Kr., and P.D.P.P.; Data interpretation, writing: K.L., M.K.S., T.A.O., D.M.G., J.P.T., J.M.S., J. C.-C., D.L., D.J.T., E.L.G., W.Z., I.P.M.T., A.B., S.J.R., S.J.C., D.F.E., G.C.-T., S.A.G., A.B.S., and R.A.E.; Provided samples and/or genetic data: S.P.K., S.L., P.Kr., P.D.P.P., K.L., M.K.S., T.A.O., D.M.G., J.P.T., J.M.S., J. C.-C., D.L., D.J.T., E.L.G., W.Z., I.P.M.T., A.B., S.J.R., S.J.C., D.F.E., G.C.-T., S.A.G., A.B.S., R.A.E., A.G.M.A., F.M.A., A. B.-F., L.B., C.B., H.B., S.B., D.C., M.C., C.C., Z.C., M.B.D., A.duB., A.B.E., A.E., P.F., J.M.F., J.G., G.G.G., R.J.Ha., H.R.H., F.H., M.H., P.H., R.-Y.H., L.I., A.I., An.J. Al.J., E.M.J., P.Ka., B.Y.K., E.K., L.A.K., S.K.K., R.K., P.Kl., M.K., B.K., C.K., D.L., A.L., T.M., A.D.M., A.N.M., K.Mo., K.Mu., S.F.N., K.O., H.O., T.-W.P.-S., J.B.P., P.P., A.P., R.P., P.R., H.A.R., I.B.R., I.K.R., R.J.S., V.W.S., N.S., W.S., B.S., L.M. S., C.L.T., L.J.T., C.T., N.U., A.M.vanA., A.V.-G., I.V., R.A.V., J.V., S.J.W., R.W., H.Y., and the PRACTICAL consortium, CRUK, BPC3, CAPS, PEGASUS; Critical review and approval of manuscript: All authors.

## For the breast cancer GWAS data

*The breast cancer genome-wide association analyses were supported by the Government of Canada through Genome Canada and the Canadian Institutes of Health Research, the ‘Ministère de l’Économie, de la Science et de l’Innovation du Québec’ through Genome Québec and grant PSR-SIIRI-701, The National Institutes of Health (U19 CA148065, X01HG007492), Cancer Research UK (C1287/A10118, C1287/A16563, C1287/A10710) and The European Union (HEALTH-F2-2009-223175 and H2020 633784 and 634935). All studies and funders are listed in Michailidou et al (2017)*.

## For the prostate cancer GWAS data

*The Prostate cancer genome-wide association analyses are supported by the Canadian Institutes of Health Research, European Commission’s Seventh Framework Programme grant agreement n° 223175 (HEALTH-F2-2009-223175), Cancer Research UK Grants C5047/A7357, C1287/A10118, C1287/A16563, C5047/A3354, C5047/A10692, C16913/A6135, and The National Institute of Health (NIH) Cancer Post-Cancer GWAS initiative grant: No. 1 U19 CA 148537-01 (the GAME-ON initiative)*.

*We would also like to thank the following for funding support: The Institute of Cancer Research and The Everyman Campaign, The Prostate Cancer Research Foundation, Prostate Research Campaign UK (now PCUK), The Orchid Cancer Appeal, Rosetrees Trust, The National Cancer Research Network UK, The National Cancer Research Institute (NCRI) UK. We are grateful for support of NIHR funding to the NIHR Biomedical Research Centre at The Institute of Cancer Research and The Royal Marsden NHS Foundation Trust*.

*The Prostate Cancer Program of Cancer Council Victoria also acknowledge grant support from The National Health and Medical Research Council, Australia (126402, 209057, 251533*,, *396414, 450104, 504700, 504702, 504715, 623204, 940394, 614296,), VicHealth, Cancer Council Victoria, The Prostate Cancer Foundation of Australia, The Whitten Foundation, PricewaterhouseCoopers, and Tattersall’s. EAO, DMK, and EMK acknowledge the Intramural Program of the National Human Genome Research Institute for their support. We also acknowledge Prostate Cancer Canada and funding by the Movember Foundation – Grant # D2013-36*.

*Genotyping of the OncoArray was funded by the US National Institutes of Health (NIH) [U19 CA 148537 for ELucidating Loci Involved in Prostate cancer SuscEptibility (ELLIPSE) project and X01HG007492 to the Center for Inherited Disease Research (CIDR) under contract number HHSN268201200008I] and by Cancer Research UK grant A8197/A16565. Additional analytic support was provided by NIH NCI U01 CA188392 (PI: Schumacher)*.

*Funding for the iCOGS infrastructure came from: the European Community’s Seventh Framework Programme under grant agreement n° 223175 (HEALTH-F2-2009-223175) (COGS), Cancer Research UK (C1287/A10118, C1287/A 10710, C12292/A11174, C1281/A12014, C5047/A8384, C5047/A15007, C5047/A10692, C8197/A16565), the National Institutes of Health (CA128978) and Post-Cancer GWAS initiative (1U19 CA148537, 1U19 CA148065 and 1U19 CA148112 – the GAME-ON initiative), the Department of Defence (W81XWH-10-1-0341), the Canadian Institutes of Health Research (CIHR) for the CIHR Team in Familial Risks of Breast Cancer, Komen Foundation for the Cure, the Breast Cancer Research Foundation, and the Ovarian Cancer Research Fund*.

*The BPC3 was supported by the U*.*S. National Institutes of Health, National Cancer Institute (cooperative agreements U01-CA98233 to D*.*J*.*H*., *U01-CA98710 to S*.*M*.*G*., *U01-CA98216 toE*.*R*., *and U01-CA98758 to B*.*E*.*H*., *and Intramural Research Program of NIH/National Cancer Institute, Division of Cancer Epidemiology and Genetics)*.

*CAPS GWAS study was supported by the Swedish Cancer Foundation (grant no 09-0677, 11-484, 12-823), the Cancer Risk Prediction Center (CRisP), a Linneus Centre (Contract ID 70867902) financed by the Swedish Research Council, Swedish Research Council (grant no K2010-70X-20430-04-3, 2014-2269)*

*PEGASUS was supported by the Intramural Research Program, Division of Cancer Epidemiology and Genetics, National Cancer Institute, National Institutes of Health*.

*The coordination of EPIC was financially supported by the European Commission (DG-SANCO) and the International Agency for Research on Cancer. The national cohorts (that recruited male participants) are supported by Danish Cancer Society (Denmark); German Cancer Aid, German Cancer Research Center (DKFZ), Federal Ministry of Education and Research (BMBF), Deutsche Krebshilfe, Deutsches Krebsforschungszentrum and Federal Ministry of Education and Research (Germany); the Hellenic Health Foundation (Greece); Associazione Italiana per la Ricerca sul Cancro-AIRC-Italy and National Research Council (Italy); Dutch Ministry of Public Health, Welfare and Sports (VWS), Netherlands Cancer Registry (NKR), LK Research Funds, Dutch Prevention Funds, Dutch ZON, World Cancer Research Fund (WCRF), Statistics Netherlands (The Netherlands); Health Research Fund (FIS), PI13/00061 to Granada;, PI13/01162 to EPIC-Murcia), Regional Governments of Andalucía, Asturias, Basque Country, Murcia and Navarra, ISCIII RETIC (RD06/0020) (Spain); Swedish Cancer Society, Swedish Research Council and County Councils of Skåne and Västerbotten (Sweden); Cancer Research UK (14136 to EPIC-Norfolk; C570/A16491 and C8221/A19170 and A29017 to EPIC-Oxford), Medical Research Council (1000143 to EPIC-Norfolk, MR/M012190/1 to EPIC-Oxford) (United Kingdom)*.

## For the ovarian cancer GWAS data

*The ovarian cancer genome-wide association analyses were supported by the U*.*S. National Institutes of Health (CA1×01HG007491-01 (C*.*I. Amos), U19-CA148112 (T*.*A. Sellers), R01-CA149429 (C*.*M. Phelan) R01-CA058598 (M*.*T. Goodman) and R01-CA211707, R01-CA204954, and R01-CA211575 (S*.*A. Gayther); Canadian Institutes of Health Research (MOP-86727 (L*.*E. Kelemen)) and the Ovarian Cancer Research Fund (A. Berchuck). The COGS project was funded through a European Commission’s Seventh Framework Programme grant (agreement number 223175 – HEALTH-F2–2009-223175). The Gynaecological Oncology Biobank at Westmead, a member of the Australasian Biospecimen Network-Oncology group, was funded by the National Health and Medical Research Council Enabling Grants ID 310670 & ID 628903 and the Cancer Institute NSW Grants ID 12/RIG/1-17 & 15/RIG/1-16. All studies and funders are listed in Phelan et al (2017)*.

## For the endometrial cancer GWAS data

*The endometrial cancer genome-wide association analyses were supported by the National Health and Medical Research Council of Australia (APP552402, APP1031333, APP1109286, APP1111246 and APP1061779), the U*.*S. National Institutes of Health (R01-CA134958), European Research Council (EU FP7 Grant), Wellcome Trust Centre for Human Genetics (090532/Z/09Z) and Cancer Research UK. OncoArray genotyping of ECAC cases was performed with the generous assistance of the Ovarian Cancer Association Consortium (OCAC), which was funded through grants from the U*.*S. National Institutes of Health (CA1×01HG007491-01 (C*.*I. Amos), U19-CA148112 (T*.*A. Sellers), R01-CA149429 (C*.*M. Phelan) and R01-CA058598 (M*.*T. Goodman); Canadian Institutes of Health Research (MOP-86727 (L*.*E. Kelemen)) and the Ovarian Cancer Research Fund (A. Berchuck). We particularly thank the efforts of Cathy Phelan. OncoArray genotyping of the BCAC controls was funded by Genome Canada Grant GPH-129344, NIH Grant U19 CA148065, and Cancer UK Grant C1287/A16563. All studies and funders are listed in O’Mara et al (2018)*.

