## Supplementary Figures 1-2 and Notes 1-2 for "Combining genome-wide studies of breast, prostate, ovarian and endometrial cancers maps cross-cancer susceptibility loci and identifies new genetic associations"

**Supplementary Figure 1. Quantile-quantile plots from the (a) main and the (b) subtype-focused meta-analyses.** Negative logarithm (base 10) *P*-values from each meta-analysis are plotted on the Y-axis. For **(a)** $\lambda$ = 1.11 and $\lambda_{1000}$ = 1.00. For **(b)** $\lambda$ = 1.15 and $\lambda_{1000}$ = 1.00. Lambda, $\lambda$, is the genomic control inflation statistic and $\lambda_{1000}$ is the same statistic scaled for 1,000 cases and 1,000 controls.

**a**

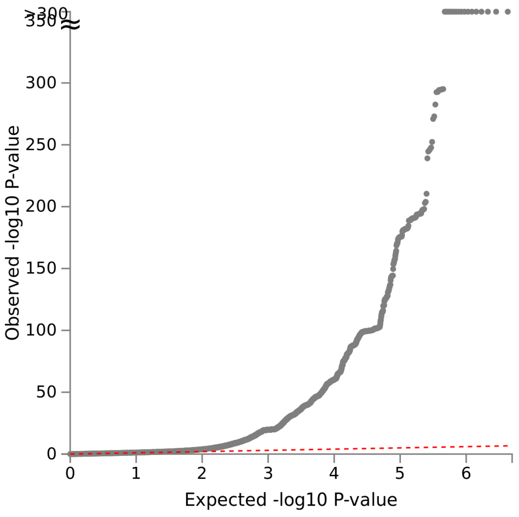


**b**

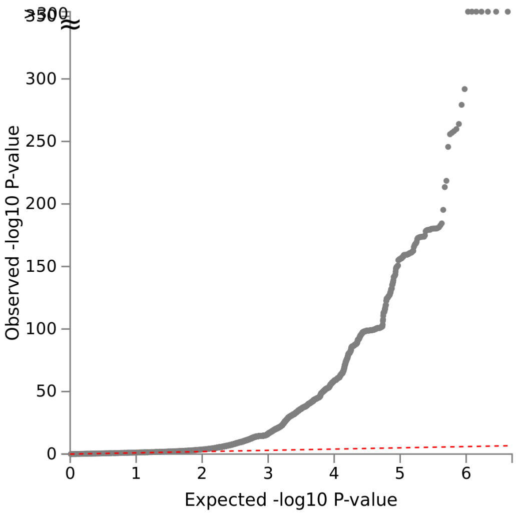


**Supplementary Figure 2. Manhattan plots of genome-wide association results from the (a) main and the (b) subtype-focused meta-analyses.** Negative logarithm (base 10) *P*-values from the meta-analysis are plotted on the Y-axis. The red dashed line indicates genome-wide significance ($P < 5 \times 10^{-8}$).

**a**

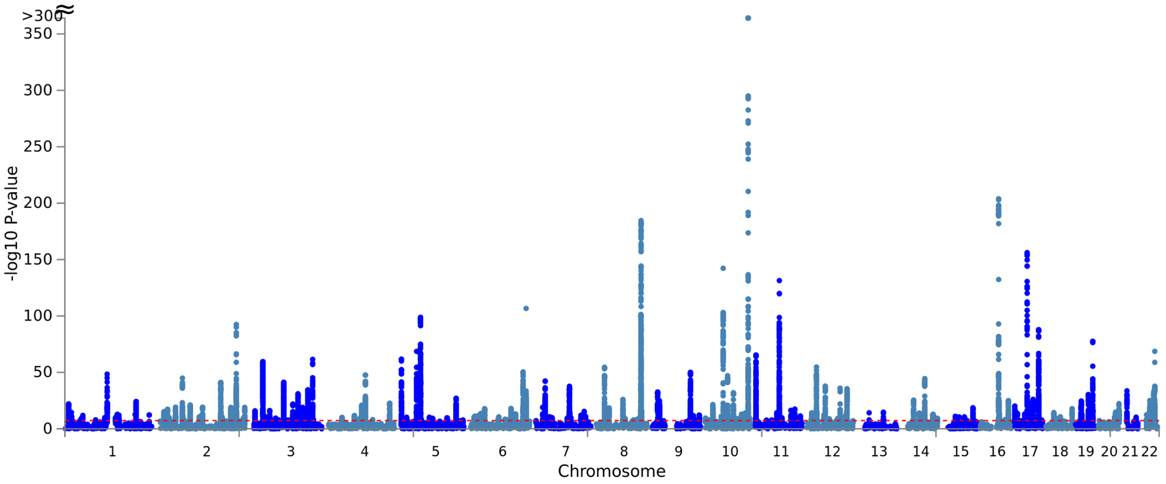


**b**

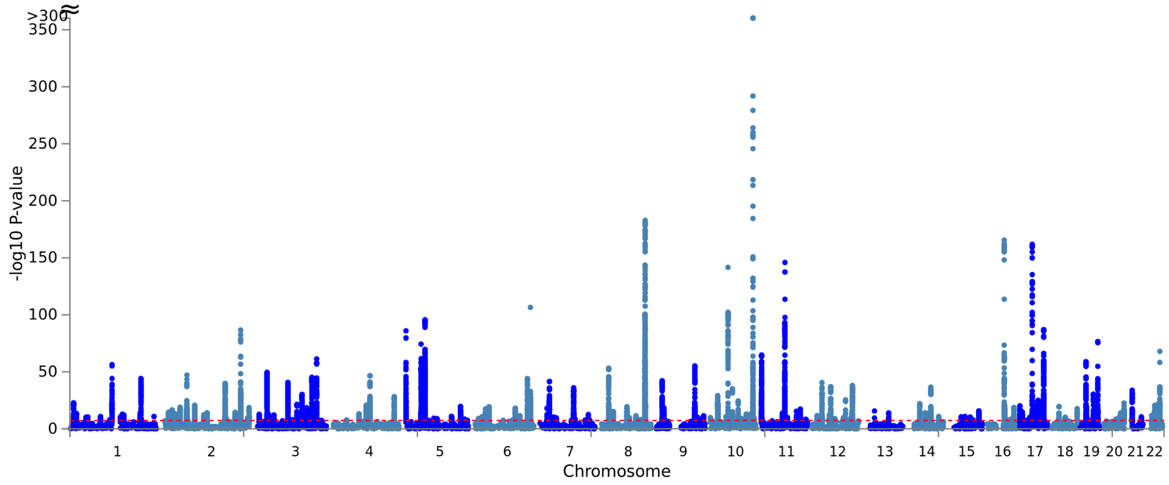


**Supplementary Note 1**

Gene-level MAGMA association analyses also revealed candidate multi-cancer susceptibility genes likely acting via diverse but fundamental carcinogenic processes. SNPs in or near *BSN* are highly pleiotropic and known to be associated at genome-wide significance ($P < 5 \times 10^{-8}$) with circulating levels of the inflammatory marker, C-reactive protein[1], and with multiple chronic inflammatory diseases[2]. *SIK2* encodes a centrosome kinase essential for bipolar mitotic spindle formation[3]. The kinase positively regulates cell-cycle progression and negatively regulates the proto-oncogene *CREB1* in prostate cancer cells[4]. *INTS10* codes for a component of Integrator, a complex involved in the transcription and processing of the small nuclear RNAs *U1* and *U2*[5]. Recurrent somatic mutations in *U1*, which regulates alternative splicing of cancer driver genes, have recently been identified in multiple tumor types[6]. *UQCC1* polymorphisms are associated at genome-wide significance with non-melanoma skin cancer[7]. *CLIC6* encodes a chloride intracellular channel and its expression has been shown to differ by *TP53* mutation and ER status in breast tumors[8]. *ATP5B* encodes a subunit of mitochondrial ATP synthase, the core enzyme of oxidative phosphorylation. The enzyme is downregulated across cancers[9], a reflection of the "Warburg effect" wherein cancer cells, regardless of their tissue of origin, favour glycolysis over oxidative phosphorylation even under aerobic conditions[10]. *IKZF2* encodes the hematopoietic-specific transcription factor "Helios" that is involved in the regulation of lymphocyte development. Helios-dependent regulation of gene expression and lymphocyte reprogramming is particularly observed in the tumor microenvironment[11]. *PYGB* encodes brain-type glycogen phosphorylase that helps metabolize glycogen as an energy source under hypoxic conditions. *PYGB* knockdown has been shown to reduce the wound-healing capability of MCF-7 breast cancer cells and the invasive potential of MDA-MB-231 breast cancer cells[12]. *PYGB* silencing in PC3 prostate cancer cells supresses growth and promotes apoptosis via NF-κB/Nrf2 signaling[13]. *PRDM5*, which encodes an epigenetic modifier that acts as a tumor suppressor, has a CpG island promoter that is silenced in multiple tumor types[14,15]. Inherited genetic variation in *LAMC1* is associated at genome-wide significance with colorectal cancer[16] and with the hormone-related traits of acne[17] and male-pattern baldness[18]. *GLIS1* was identified in our study based on its association with breast and prostate cancer. GLIS1 is a hypoxia-inducible transcription factor that regulates gene expression related to cell migration and invasion in breast cancer cells, specifically transcriptional activation of the Wnt/β-catenin pathway[19,20]. *CDKN1A* and *TIAL1* were identified at gene-level genome-wide significance ($P_{\mathrm{MAGMA}} < 2.62 \times 10^{-6}$) for breast cancer alone and in the combined data, with the breast cancer association underlying the combined signal. Variants in/near these genes are associated at sub-genome-wide significance levels ($5 \times 10^{-8} < P_{\mathrm{lead\ SNP}} \leq 10^{-7}$), with breast cancer susceptibility in the Michailidou et al. data set.[21] *TIAL1* encodes nucleolysin TIAR that has roles in G1/S phase cell-cycle arrest and caspase-dependent apoptosis[22]. *CDKN1A* expression is tightly controlled by TP53 and mediates the G1/S phase arrest in response to a range of stress stimuli[23]. These genes add to a list of candidate breast cancer susceptibility genes identified by GWAS, including *CCND1*[24] and *CCNE1*[25], that collectively underscore the importance of the G1/S checkpoint as a major functional effector of inherited breast cancer risk.

**Supplementary Note 2**


### PIs from the PRACTICAL (http://practical.icr.ac.uk/), CRUK, BPC3, CAPS, PEGASUS consortia:

Rosalind A. Eeles[1,2], Christopher A. Haiman[3], ZSofia Kote-Jarai[1], Fredrick R. Schumacher[4,5], Sara Benlloch[6,1], Ali Amin Al Olama[6,9], Kenneth Muir[7,8], Sonja I. Berndt[10], David V. Conti[3], Fredrik Wiklund[11], Stephen Chanock[10], Susan M. Gapstur[12], Victoria L. Stevens[12], Catherine M. Tangen[13], Jyotsna Batra[15,16], Judith Clements[15,16], APCB BioResource[15], Henrik Gronberg[11], Nora Pashayan[17,18], Johanna Schleutker[19,20], Demetrius Albanes[21], Stephanie Weinstein[21], Alicja Wolk[22, 23], Catharine West[24], Lorelei Mucci[25], Géraldine Cancel-Tassin[26,27], Stella Koutros[10], Karina Dalsgaard Sorensen[28,29], Eli Marie Grindedal[30], David E. Neal[31,32,33], Freddie C. Hamdy[34,35], Jenny L. Donovan[36], Ruth C. Travis[37], Robert J. Hamilton[38,39], Sue Ann Ingles[3], Barry S. Rosenstein[40], Yong-Jie Lu[41], Graham G. Giles[42,43,44], Adam S. Kibel[45], Ana Vega[46,47,48], Manolis Kogevinas[49,50,51,52], Kathryn L. Penney[53], Jong Y. Park[54], Janet L. Stanford[55,56], Cezary Cybulski[57], Børge G. Nordestgaard[58,59], Sune F. Nielsen[58,59], Hermann Brenner[60,61,62], Christiane Maier[63], Jeri Kim[64], Esther M. John[65], Manuel R. Teixeira[66,67], Susan L. Neuhausen[68], Kim De Ruyck[69], Azad Razack[70], Lisa F. Newcomb[55,71], Davor Lessel[72], Radka Kaneva[73], Nawaid Usmani[74,75], Frank Claessens[76], Paul A. Townsend[77], Manuela Gago-Dominguez[78,79], Monique J. Roobol[80], Florence Menegaux[81], Kay-Tee Khaw[82], Lisa Cannon-Albright[83,84], Hardev Pandha[85], Stephen N. Thibodeau[86], David J Hunter[87], Peter Kraft[87], William J. Blot[88,89], Elio Riboli[90],

1 The Institute of Cancer Research, London, UK.
2 Royal Marsden NHS Foundation Trust, London, UK.
3 Department of Preventive Medicine, Keck School of Medicine, University of Southern California/Norris Comprehensive Cancer Center, Los Angeles, CA 90015, USA
4 Department of Population and Quantitative Health Sciences, Case Western Reserve University, Cleveland, OH 44106-7219, USA
5 Seidman Cancer Center, University Hospitals, Cleveland, OH 44106, USA.
6 Centre for Cancer Genetic Epidemiology, Department of Public Health and Primary Care, University of Cambridge, Strangeways Research Laboratory, Cambridge, UK
7 Division of Population Health, Health Services Research and Primary Care, University of Manchester, Oxford Road, Manchester, M13 9PL, UK
8 Warwick Medical School, University of Warwick, Coventry, UK.
9 University of Cambridge, Department of Clinical Neurosciences, Stroke Research Group, R3, Box 83, Cambridge Biomedical Campus, Cambridge CB2 0QQ, UK
10 Division of Cancer Epidemiology and Genetics, National Cancer Institute, NIH, Bethesda, Maryland, 20892, USA
11 Department of Medical Epidemiology and Biostatistics, Karolinska Institute, Stockholm, Sweden.
12 Epidemiology Research Program, American Cancer Society, 250 Williams Street, Atlanta, GA 30303, USA
13 SWOG Statistical Center, Fred Hutchinson Cancer Research Center, Seattle, WA 98109, USA
15 Australian Prostate Cancer Research Centre-Qld, Institute of Health and Biomedical Innovation and School of Biomedical Sciences, Queensland University of Technology, Brisbane QLD 4059, Australia
16 Translational Research Institute, Brisbane, Queensland 4102, Australia
17 University College London, Department of Applied Health Research, London, UK.
18 Centre for Cancer Genetic Epidemiology, Department of Oncology, University of Cambridge, Strangeways Laboratory, Cambridge, UK.
19 Institute of Biomedicine, Kiinamyllynkatu 10, FI-20014 University of Turku, Finland

20 Department of Medical Genetics, Genomics, Laboratory Division, Turku University Hospital, PO Box 52, 20521 Turku, Finland
21 Division of Cancer Epidemiology and Genetics, National Cancer Institute, NIH, Bethesda, MD 20892, USA
22 Division of Nutritional Epidemiology, Institute of Environmental Medicine, Karolinska Institutet, SE-171 77 Stockholm, Sweden
23 Department of Surgical Sciences, Uppsala University, Uppsala, Sweden
24 Division of Cancer Sciences, University of Manchester, Manchester Academic Health Science Centre, Radiotherapy Related Research, The Christie Hospital NHS Foundation Trust, Manchester, M13 9PL UK
25 Department of Epidemiology, Harvard T.H. Chan School of Pubic Health, Boston, MA, USA.
26 CeRePP, Tenon Hospital, Paris, France
27 Sorbonne Universite, GRC n°5, AP-HP, Tenon Hospital, 4 rue de la Chine, F-75020 Paris, France
28 Department of Molecular Medicine, Aarhus University Hospital, Palle Juul-Jensen Boulevard 99, 8200 Aarhus N, Denmark
29 Department of Clinical Medicine, Aarhus University, DK-8200 Aarhus N
30 Department of Medical Genetics, Oslo University Hospital, Norway.
31 Nuffield Department of Surgical Sciences, University of Oxford, Room 6603, Level 6, John Radcliffe Hospital, Headley Way, Headington, Oxford, OX3 9DU, UK
32 University of Cambridge, Department of Oncology, Addenbrooke's Hospital, Cambridge, UK.
33 Cancer Research UK Cambridge Research Institute, Li Ka Shing Centre, Cambridge, UK.
34 Nuffield Department of Surgical Sciences, University of Oxford, Oxford, OX1 2JD, UK
35 Faculty of Medical Science, University of Oxford, John Radcliffe Hospital, Oxford, UK
36 Population Health Sciences, Bristol Medical School, University of Bristol, BS8 2PS, UK
37 Cancer Epidemiology Unit, Nuffield Department of Population Health University of Oxford, Oxford, UK.
38 Dept. of Surgical Oncology, Princess Margaret Cancer Centre, Toronto, Canada.
39 Dept. of Surgery (Urology), University of Toronto, Canada
40 Department of Radiation Oncology and Department of Genetics and Genomic Sciences, Box 1236, Icahn School of Medicine at Mount Sinai, One Gustave L. Levy Place, New York, NY 10029, USA
41 Centre for Molecular Oncology, Barts Cancer Institute, Queen Mary University of London, John Vane Science Centre, Charterhouse Square, London, EC1M 6BQ, UK
42 Cancer Epidemiology Division, Cancer Council Victoria, 615 St Kilda Road, Melbourne, Victoria 3004, Australia.
43 Centre for Epidemiology and Biostatistics, Melbourne School of Population and Global Health, The University of Melbourne, Grattan Street, Parkville, VIC 3010, Australia
44 Precision Medicine, School of Clinical Sciences at Monash Health. Monash University. Clayton, Victoria, Australia, 3168
45 Division of Urologic Surgery, Brigham and Womens Hospital, 75 Francis Street, Boston, MA 02115, USA
46 Fundación Pública Galega de Medicina Xenómica, Santiago de Compostela, 15706, Spain.
47 Instituto de Investigación Sanitaria de Santiago de Compostela, Santiago De Compostela, 15706, Spain
48 Centro de Investigación en Red de Enfermedades Raras (CIBERER), Spain
49 ISGlobal, Barcelona, Spain.
50 IMIM (Hospital del Mar Research Institute), Barcelona, Spain.
51 Universitat Pompeu Fabra (UPF), Barcelona, Spain.
52 CIBER Epidemiologia y Salud Publica (CIBERESP), Madrid, Spain

53 Channing Division of Network Medicine, Department of Medicine, Brigham and Women's Hospital/Harvard Medical School, Boston, MA, USA.
54 Department of Cancer Epidemiology, Moffitt Cancer Center, 12902 Magnolia Drive, Tampa, FL 33612, USA
55 Division of Public Health Sciences, Fred Hutchinson Cancer Research Center, Seattle, Washington, 98109-1024, USA
56 Department of Epidemiology, School of Public Health, University of Washington, Seattle, Washington, USA.
57 International Hereditary Cancer Center, Department of Genetics and Pathology, Pomeranian Medical University, Szczecin, Poland.
58 Faculty of Health and Medical Sciences, University of Copenhagen, Denmark.
59 Department of Clinical Biochemistry, Herlev and Gentofte Hospital, Copenhagen University Hospital, Herlev, Denmark.
60 Division of Clinical Epidemiology and Aging Research, German Cancer Research Center (DKFZ), Heidelberg, Germany.
61 German Cancer Consortium (DKTK), German Cancer Research Center (DKFZ), Heidelberg, Germany.
62 Division of Preventive Oncology, German Cancer Research Center (DKFZ) and National Center for Tumor Diseases (NCT), Heidelberg, Germany.
63 Institute for Human Genetics, University Hospital Ulm, Ulm, Germany.
64 The University of Texas M. D. Anderson Cancer Center, Department of Genitourinary Medical Oncology, Houston, TX, USA.
65 Department of Medicine, Division of Oncology, Stanford Cancer Institute, Stanford University School of Medicine, Stanford, 780 Welch Road, CJ250C, CA 94304-5769
66 Department of Genetics, Portuguese Oncology Institute of Porto, Porto, Portugal.
67 Biomedical Sciences Institute (ICBAS), University of Porto, Porto, Portugal.
68 Department of Population Sciences, Beckman Research Institute of the City of Hope, Duarte, CA, USA.
69 Ghent University, Faculty of Medicine and Health Sciences, Basic Medical Sciences, Gent, Belgium.
70 Department of Surgery, Faculty of Medicine, University of Malaya, Kuala Lumpur, Malaysia.
71 Department of Urology, University of Washington, 1959 NE Pacific Street, Box 356510, Seattle, WA 98195, USA
72 Institute of Human Genetics, University Medical Center Hamburg-Eppendorf, Hamburg, Germany.
73 Molecular Medicine Center, Department of Medical Chemistry and Biochemistry, Medical University, Sofia, Bulgaria.
74 Department of Oncology, Cross Cancer Institute, University of Alberta, Edmonton, Alberta, Canada.
75 Division of Radiation Oncology, Cross Cancer Institute, Edmonton, Alberta, Canada.
76 Molecular Endocrinology Laboratory, Department of Cellular and Molecular Medicine, KU Leuven, Leuven, Belgium.
77 Division of Cancer Sciences, Manchester Cancer Research Centre, Faculty of Biology, Medicine and Health, Manchester Academic Health Science Centre, NIHR Manchester Biomedical Research Centre, Health Innovation Manchester, Univeristy of Manchester, UK.
78 Genomic Medicine Group, Galician Foundation of Genomic Medicine, Instituto de Investigacion Sanitaria de Santiago de Compostela (IDIS), Complejo Hospitalario Universitario de Santiago, Servicio Galego de Saúde, SERGAS, Santiago De Compostela, Spain.
79 University of California San Diego, Moores Cancer Center, La Jolla, CA, USA.
80 Department of Urology, Erasmus University Medical Center, Rotterdam, the Netherlands.

81 Université Paris-Saclay, UVSQ, Inserm, CESP, Villejuif, France.
82 Clinical Gerontology Unit, University of Cambridge, Cambridge, UK.
83 Division of Genetic Epidemiology, Department of Medicine, University of Utah School of Medicine, Salt Lake City, Utah, USA.
84 George E. Wahlen Department of Veterans Affairs Medical Center, Salt Lake City, UT, USA.
85 The University of Surrey, Guildford, Surrey, UK.
86 Department of Laboratory Medicine and Pathology, Mayo Clinic, Rochester, MN, USA.
87 Program in Genetic Epidemiology and Statistical Genetics, Department of Epidemiology, Harvard T.H. Chan School of Public Health, Boston, MA, USA.
88 Division of Epidemiology, Department of Medicine, Vanderbilt University Medical Center, TN, USA.
89 International Epidemiology Institute, Rockville, MD, USA
90 Department of Epidemiology and Biostatistics, School of Public Health, Imperial College London, SW7 2AZ, UK
